# Phylogenetically distant enzymes localized in cytosol and plastids drive citral biosynthesis in lemongrass

**DOI:** 10.1101/2024.03.11.583845

**Authors:** Priyanka Gupta, Anuj Sharma, N.R. Kiran, T.K. Pranav Raj, Ram Krishna, Dinesh A. Nagegowda

## Abstract

Except for the genetic basis of citral-forming alcohol dehydrogenases (ADHs) in *Litsea cubeba* tree, and biochemical studies on citral-forming enzymes from select plants, knowledge regarding *in-planta* biosynthesis of citral and its metabolic origin remains limited. Here, we have elucidated the functions of an ADH (CfADH1) and an aldoketo-reductase (CfAKR2b) in citral biosynthesis in lemongrass (*Cymbopogon flexuosus*), one of the most cultivated aromatic crops for its citral-rich essential oil. Expression of both *CfADH1* and *CfAKR2b* showed correlation with citral accumulation in different developmental stages. Recombinant CfADH1 and CfAKR2b, despite their sequence unrelatedness, exhibited similar kinetic properties and formed citral from geraniol with NADP cofactor. Virus-induced gene silencing in lemongrass, and transient expression in lemon balm (*Melissa officinalis*), demonstrated the *in-planta* involvement of *CfADH1* and *CfAKR2b* in citral biosynthesis. While CfADH1 exhibited a dual cytosolic/plastidial localization, CfAKR2b was localized to cytosol. Moreover, feeding lemongrass seedlings with mevalonate- and methylerythritol-phosphate-pathway specific inhibitors combined with volatile profiling supported the role of both pathways in citral formation. Our results demonstrate phylogenetically distant enzymes localized in cytosol and plastids drive citral biosynthesis in lemongrass, indicating an evolutionary scenario aimed at maximizing the utilization of precursor pools from both cytosolic and plastidial pathways for high citral production.

## Introduction

Plants continue to amaze us with their intricate mechanisms for producing terpenoids. Initially, it was widely believed that all terpenes originated from the cytoplasm-localized MVA pathway. However, the discovery of the plastid-localized MEP pathway revolutionized our understanding of terpenoid biosynthesis (Philips *et al*., 2008). Accordingly, it has been established that terpenes are derived from the universal five-carbon precursors, isopentenyl diphosphate (IPP) and dimethylallyl diphosphate (DMAPP), which are formed via either the cytosolic mevalonic acid (MVA) pathway or the plastidial methylerythritol phosphate (MEP) pathway (Nagegowda, 2010). Monoterpenes are produced from the C_10_ prenyl diphosphate precursor geranyl diphosphate (GPP), which is typically formed by the condensation of plastidial IPP and DMAPP pool through the activity of geranyl diphosphate synthase (GPPS). The transformation of geranyl diphosphate (GPP) into monoterpenes is facilitated by the action of terpene synthases belonging to monoterpene synthase class (Nagegowda and Gupta, 2020). With advancements in research, new, non-canonical pathways of terpenoid biosynthesis are being uncovered, expanding our comprehension of the remarkable diversity and complexity within plant terpenoid biochemistry. For instance, the traditional belief that monoterpenes are exclusively derived from the plastidial MEP pathway has been challenged by several studies, which have shown that monoterpene biosynthesis can also occur in the cytosol (Ohara, 2003; Aharoni *et al*., 2004; Wu *et al*., 2006; Davidovich-Rikanati *et al*., 2008; Dong *et al*., 2013, 2016) Moreover, recent studies in Rose and few other plants have demonstrated that a non-canonical TPS-independent and cytosolic route for the formation of monoterpene, geraniol (Bergman *et al*., 2021b; Conart *et al*., 2022).

Citral (3,7-Dimethyl-2,6-Octadienal) is a naturally occurring acyclic monoterpene aldehyde. It imparts a refreshing and zesty scent reminiscent of citrus fruits and is composed of a mixture of two stereoisomers: geranial (*trans*-citral or α-citral) and neral (*cis*-citral or β-citral). Due to its strong lemony essence, citral is utilized in the fragrance industry to create citrus-scented perfumes, colognes, and household products (Sharmeen *et al*., 2021). It is also used as a precursor for the production of ionone and vitamin A (Parker *et al*., 2016). Citral is reported to be found in various plants but only few plants produce it in abundance. Its formation in plants has been reported to involve the enzymatic oxidation of geraniol and or nerol to citral by enzymes belonging to NADP-dependent alcohol dehydrogenases (ADHs) class. So far, ADHs catalyzing the conversion of geraniol to citral by removing the hydrogen-oxygen bonds from geraniol and nerol have been biochemically characterized in few dicot species that include *ObADH* from sweet basil (*Ocimum basilicum*), *ADH* from *Perilla* spp., and *PmADH* from pygmy smartweed (*Persicaria minor*), and a monocot ginger ZoGeDH (Iijima *et al*., 2006; Lüddeke *et al*., 2012; Iijima *et al*., 2014; Sato-Masumoto & Ito, 2014; Tan *et al*., 2018). Besides these ADHs, another type of phylogenetically distant ADH belonging to aldoketo reductase (AKR) superfamily of NAD(P)H-dependent oxidoreductases, has been identified in *Perilla* in which the recombinant proteins were biochemically characterized (Sato-Masumoto & Ito, 2014). Similar to ADHs, these AKRs catalyzed the NADP-dependent conversion of geraniol into citral under *in vitro* conditions. However, the genetic basis for any of the above genes encoding ADHs and AKR for their involvement in citral biosynthesis has not been investigated. In contrast to the above mentioned biochemical studies, a recent report has demonstrated both biochemical and genetic basis of ADH in citral formation in mountain pepper (*Litsea cubeba*), a member of the Lauraceae family. This study revealed the involvement of two ADHs, LcADH28 and LcADH29, are responsible for catalyzing the conversion of geraniol to geranial (α-citral) and nerol to neral (β-citral), respectively (Zhao *et al*., 2024). Additionally, the metabolic origin of citral remains unclear in plants, though some earlier studies involving carbon isotope feeding in *Cymbopogon* have indicated the potential involvement of MVA and MEP pathways in geraniol and citral formation, respectively (Akhila, 1985; Gupta & Ganjewala, 2015).

The ability to produce citral is widespread in dicots, but among monocots only a few species, including aromatic grasses of the *Cymbopogon* genus, have evolved this ability. Lemongrass (*Cymbopogon flexuosus*), is a perennial herb and one of the commercially important aromatic grasses belonging to Poaceae family, due to its citral-rich (around 70-80%) essential oil, making it the primary source of citral (Kumar *et al*., 2023). Besides citral, the lemongrass essential oil also contains minor amounts of other monoterpenes such as geraniol, citronellal, and myrcene (Mukarram *et al*., 2021). Our previous study in lemongrass and a recent study by other researchers in different *Cymbopogon* spp, involving transcriptomics, gene expression analyses, and molecular modelling, showed evidence for the presence of ADH and AKR encoding genes, indicating the potential involvement of any of these enzymes in citral biosynthesis (Meena *et al*., 2016; Bhat *et al*., 2023). Here, we have explored the biosynthesis of citral and its origin in lemongrass using a combination of biochemical, reverse and forward genetics, and molecular approaches. Our results for the first time reveal that phylogenetically distant ADH (CfADH1) and AKR (CfAKR2b) enzymes localized in the cytosol and plastids play integral roles in lemongrass citral biogenesis.

## Materials and Methods

### Plant material and tissue collection

*Cymbopogon flexuosus* cv. Krishna (National Gene Bank, CSIR-CIMAP, Lucknow, India) plants were used for all experiments in the study. For enzyme assay and volatile collection, leaves from mature plants grown in field conditions were used. For determining stage-specific expression and volatile analyses, leaves from different developmental stages (1 month, 2 month, 3 month and harvesting stage) were collected and used for RNA extraction and GC analysis. For cytosolic and plastidial fractionation, young leaves from field grown plants were collected and used.

### Phylogenetic analysis

For phylogenetic analysis, amino acid sequences of characterized ADH and AKR proteins from different plant species were retrieved from the National Center for Biotechnology Information (NCBI) data base (www.ncbi.nlm.nih.gov). The phylogentic tree was constructed by neighbour-joining method using MEGA11 software (Tamura *et al*., 2021) Multiple sequence alignment of proteins was performed using ClustalW (Thompson *et al*., 1994) with default parameters through https://www.genome.jp/tools-bin/clustalw.

### Generation of silencing and overexpression constructs

The pTRV1 and pTRV2 vectors used for generating VIGS constructs were procured from The Arabidopsis Information Resource (TAIR), USA. The 551 bp, 570 bp, and 471 bp fragments of *CfPDS, CfADH1,* and *CfAKR2b*, respectively were amplified from leaf cDNA by PCR using gene specific primers having *Eco*RI restriction site at 5′ end of each primer for *CfADH1* and *CfAKR2b*, and *Bam*HI for *CfPDS* (Table S1). The resulting PCR product was purified and sub-cloned into pJET1.2/vector and sequences were confirmed by nucleotide sequencing. The confirmed fragment was then cloned into pTRV2 vector digested with respective restriction enzymes. The resulting pTRV2::*CfPDS,* pTRV2::*CfADH1,* and pTRV2::*CfAKR2b* plasmids were confirmed by restriction digestion. For generation of overexpression constructs, the open reading frame of *CfADH1,* and *CfAKR2b* were PCR amplified using lemongrass leaf cDNA with specific forward and reverse primers (Table S1). The amplicons were cloned into pJET1.2/cloning vector for sequence confirmation, and then sub-cloned into *Nde*I and *Eco*RI sites of pRI101-AN binary vector under the control of the *35S* promoter of Cauliflower mosaic virus (CaMV) to form pRI::*CfADH1* and pRI::*CfAKR2b* constructs (Fig. S1). The VIGS and overexpression constructs were then mobilized into *Agrobacterium tumefaciens* GV3101 competent cells by freeze thaw method.

### Silencing of *CfADH1* and *CfAKR2b* in lemongrass through VIGS

A new method for VIGS was developed for lemongrass. The overnight grown *Agrobacteria* cultures harboring pTRV1 and pTRV2::*CfPDS* or pTRV2::*CfADH1* or and pTRV2::*CfAKR2b* or pTRV2 (EV) constructs were resuspended in MES buffer (MES, MgCl_2_ and Acetosyringone) and mixed in 1:1 ratio and kept in a shaker incubator for 3 h. The suspension was used to infect by injecting about 100 µl of agro-suspension at the tiller-base region of uprooted seedlings. The infected seedlings were transplanted back into pots with soilrite in a box, covered with klin-film having pores to maintain the humidity, and kept in dark for 48 h. The pots containing seedlings were removed from the box and kept in a growth room at 22 °C, 16-8h light cycle. Leaves from pTRV2::*CfPDS,* pTRV2::*CfADH1*, pTRV2::*CfAKR2b* and pTRV2 infected plants were harvested 4 weeks post infection, when albino phenotype developed in the newly emerging leaves in pTRV2::*CfPDS* infected plants. The collected tissues were divided into two parts with one part used for immediate volatile extraction and the remaining part stored in −80°C for further use for RNA extraction.

### Transient expression of *CfADH1* and *CfAKR2b* in lemon balm

Transient expression was carried out using 6-8 leaf staged lemon balm plants. *Agrobacteria* harboring p19 (suppressor of RNA silencing) and pRI::*CfADH1* or pRI::*CfAKR2b* or pRI (control) construct and were mixed in 1:1 ratio and kept in a shaker incubator for 3 h at 28°C. Agroinfiltration was performed in a vacuum desiccator by immersing the upper part of the lemon balm seedling with all the leaves into *Agrobacteria* solution in an inverted manner and vacuum was applied for 10-15 min. After infiltration, the seedlings were kept in a box and covered with klin-film having pores to maintain the humidity and kept in dark for 48 h. Next, the pots containing seedlings were removed from the box and kept in a growth room at 22 °C, 16-8h light cycle for 4 days. Leaf samples (top 4) were collected and used for volatile and RNA extraction immediately.

### RNA isolation, cDNA synthesis and qRT-PCR analysis

Total RNA was extracted from lemongrass and lemon balm leaves using the protocol by Deepa et al. (Deepa *et al*., 2014) with slight modifications. Briefly, 50mg of leaf tissue was ground to a fine powder using liquid nitrogen and the power was added with 2% polyvinylpyrrolidone (PVP) (powder form). To this 1 ml of extraction buffer was added and vortexed followed by sequential acid phenol: chloroform extraction to remove polyphenols and proteins. Further Sodium acetate is used for purification to eliminate polysaccharides. The RNA yield and purity was measured spectrophotometrically by analyzing the absorbance at A260, and monitoring the ratio at A260/280. cDNA was prepared using isolated total RNA, and qRT-PCR analysis was conducted as previously described in the literature (Singh *et al*., 2015, 2017). The cDNA was normalized using *EF1α* as an internal reference. For qRT-PCR analysis, gene-specific primers designed outside the gene region utilized for cloning into pTRV2 were employed (Table S1). The reaction was carried out using 2X SYBR green mix from Thermo Scientific (USA) and was run in the StepOne Real-Time PCR System from Applied Biosystems (USA). The qRT-PCR conditions used were as follows: 94 °C for 10 min, followed by 40 cycles of 94 °C for 15 s and 60 °C for 45 s. Fold-change differences in gene expression were analyzed using the comparative cycle threshold (*C*_t_) method.

### Protein expression and purification

The open reading frame of *CfADH1* and *CfAKR2b* was cloned into the *Nde*I/*Eco*RI sites of pET28a bacterial expression vector in-frame with the C-terminal 6X-His-tag (Novagen, http://www.emdbiosciences.com) resulting in pET28a::*CfADH1* and pET28a::*CfAKR2b,* respectively. For recombinant protein expression, pET28a::*CfADH1* and pET28a::*CfAKR2b* and pET28a (empty control) constructs were transformed into *E. coli* BL21 Rosetta-2 cells. A single colony was inoculated into 5 ml of LB with kanamycin (50 mg/L) and chloramphenicol (37 mg/L) and grown overnight at 37 °C in an incubator shaker set at 180 rpm. One ml of this culture was inoculated into 500 ml of LB with kanamycin (50 mg/L) and chloramphenicol (37 mg/L) and grown at 37 °C until the OD_600_ reached 0.4. Then the cultures were induced by the addition of 0.4 mM isopropylthio-β-galactoside (IPTG) and grown in an incubator shaker at 16 °C, 180 rpm for 16 hrs. The recombinant protein was purified by affinity chromatography using nickel-nitrilotriacetic acid (Ni-NTA) agarose beads (Qiagen, http://www.qiagen.com) as per the manufacturer’s instructions. The concentration of the purified protein was determined using the Bradford method (Bradford, 1976) and purity was determined using SDS-PAGE analysis (Fig. S2).

### Enzyme assay and kinetic analysis

Enzyme assays were performed using purified recombinant proteins of CfADH1 and CfAKR2b. The assay buffer comprised of 50 mM Tris-HCl of pH 7.5 and 1 mM DTT. Five to 10 µg each of purified CfADH1 and CfAKR2b was incubated with 1 mM substrates in a 300 µl reaction for 16 h at 30°C. The assay mix was added with 15 µl of glacial acetic acid and vortexed to stop the enzyme reaction. Next, 150 µl of n-hexane containing the internal standard (*E,E*-farnesol) was added into the assay mix, vortexed, and centrifuged to extract the volatiles. The extract was injected into GC (10 µl) or GC-MS (3 µl) for determination and quantification of volatiles. To determine the kinetic parameters, a series of dilutions for geraniol were prepared in concentrations ranging from 25 - 225 µM. When one of the substrates (in various concentrations) was added to initiate the reactions, the cofactor was kept at a constant saturating concentration (1 mM for NADP and NADPH). Product concentration was analyzed in GC using peak area as an input. Subsequently, Lineweaver–Burk double-reciprocal plot was used to derive the probable Michaelis–Menten Constant (*K*_m_) and *V*_max_ using Prism GraphPad (https://www.graphpad.com/).

### Quantification of volatiles

In all experiments, volatile terpene compounds were detected and identified using gas chromatography-mass spectrometry (GC-MS). For quantification, calibration curves were generated from known standards of the volatile compounds, and the concentration of each compound in the samples was determined using GC-MS. In addition, the concentration of volatiles in the samples was also determined using the Relative Response Factor (RRF) method in gas chromatography (GC), with *E,E*-farnesol serving as the internal standard.

### Analysis of subcellular localization

The subcellular localization of CfADH1 and CfAKR2b was determined by creating a YFP fusion protein using the pGWB441 vector. *CfADH* and *CfAKR* were cloned into pGWB441 to express them as C-terminal eYFP-tagged proteins. The resulting pGWB441::CfADH1-YFP and pGWB441::CfAKR2b-YFP constructs were mobilized into *Agrobacterium tumefaciens* (GV3101) using the freeze-thaw method. For transient expression of the recombinant proteins in *N. benthamiana* leaves, *Agrobacterium* suspension was prepared in infiltration buffer (10 mM MES, pH 5.6, 10 mM MgCl_2_, and 200 μM acetosyringone), and the agroinfiltration method was performed on 5- to 6-week-old plants. p19 (suppressor of RNA silencing), was co-expressed along with recombinant *CfADH* and *CfAKR*. After 36 to 48 hours of agroinfiltration, leaf sections were analyzed, and images were captured under a Carl Zeiss LSM880 laser scanning confocal microscope using a 63× oil immersion objective with a numerical aperture of 1.4. YFP fluorescence was excited at 514 nm and detected in the range of 525 to 562 nm, while chlorophyll autofluorescence was excited at 637 nm and detected in the range of 660 to 700 nm.

### Inhibitor feeding assay

Fosmidomycin and lovastatin were obtained from Sigma-Aldrich. Young and rapidly expanding 4-leaf stage seedlings (aged 15 days) were cut at the tiller base region and then placed in Eppendorf tubes, with one seedling per tube. Each tube contained either 500 µL of an aqueous solution of fosmidomycin (2 mM) or 500 µL of a 20% ethanolic solution of lovastatin (1 mM). An equal number of seedlings, also cut at the tiller base, were placed in tubes containing either an aqueous solution or a 20% ethanolic solution without any inhibitor, serving as controls. The tubes containing the seedlings were incubated under a 16-hour light and 8-hour dark cycle, undisturbed, for 48 hours. After incubation, the seedlings were removed from the tubes, and their leaves were used to extract volatiles using hexane containing the internal standard (*E,E*-farnesol). The extracted volatiles (geraniol and citral) were then analysed and quantified by GC analysis.

## Results

### Isolation and sequence analysis of *CfADH1* and *CfAKR2b*

Citral-forming enzymes belong to the members of the MDR (medium-chain dehydrogenases/reductases) superfamily and have only been biochemically characterized in a few species and *in planta* characterized in only *L. cubeba*. To date, there have been no reports of such enzymes in any of the aromatic grasses including lemongrass. In our prior research, the correlation between gene expression and metabolites across various *Cymbopogon* species, coupled with *in silico* modeling and docking studies involving geraniol, hinted at the involvement of both CfADH1 and CfAKR2b in citral formation in lemongrass. To further ascertain whether lemongrass leaves possess ADH activity, crude protein extracts from the leaves were incubated with geraniol and nerol, the substrate for ADH. Analysis of assay products revealed that crude protein extracts effectively converted only geraniol but not nerol into citral (Figs 1a, S3), thus indicating the involvement of ADH type of enzymes in citral formation from geraniol. Hence, to further characterize the functional role of the identified *CfADH1* and *CfAKR2b* genes in lemongrass essential oil biosynthesis, the full-length genes were cloned using cDNA derived from lemongrass leaves. The open reading frame (ORF) of CfADH1 (GenBenk Accession No: PP421063) consisted of 1254 bp, encoding a protein of 417 amino acids (aa), with a calculated molecular weight of 44.5 kDa. Whereas, the ORF of CfAKR2b (GenBenk Accession No: PP421064) encompassed 1041 bp, coding for a protein of 346 aa, with a calculated molecular weight of 38.5 kDa.

**Figure 1.**
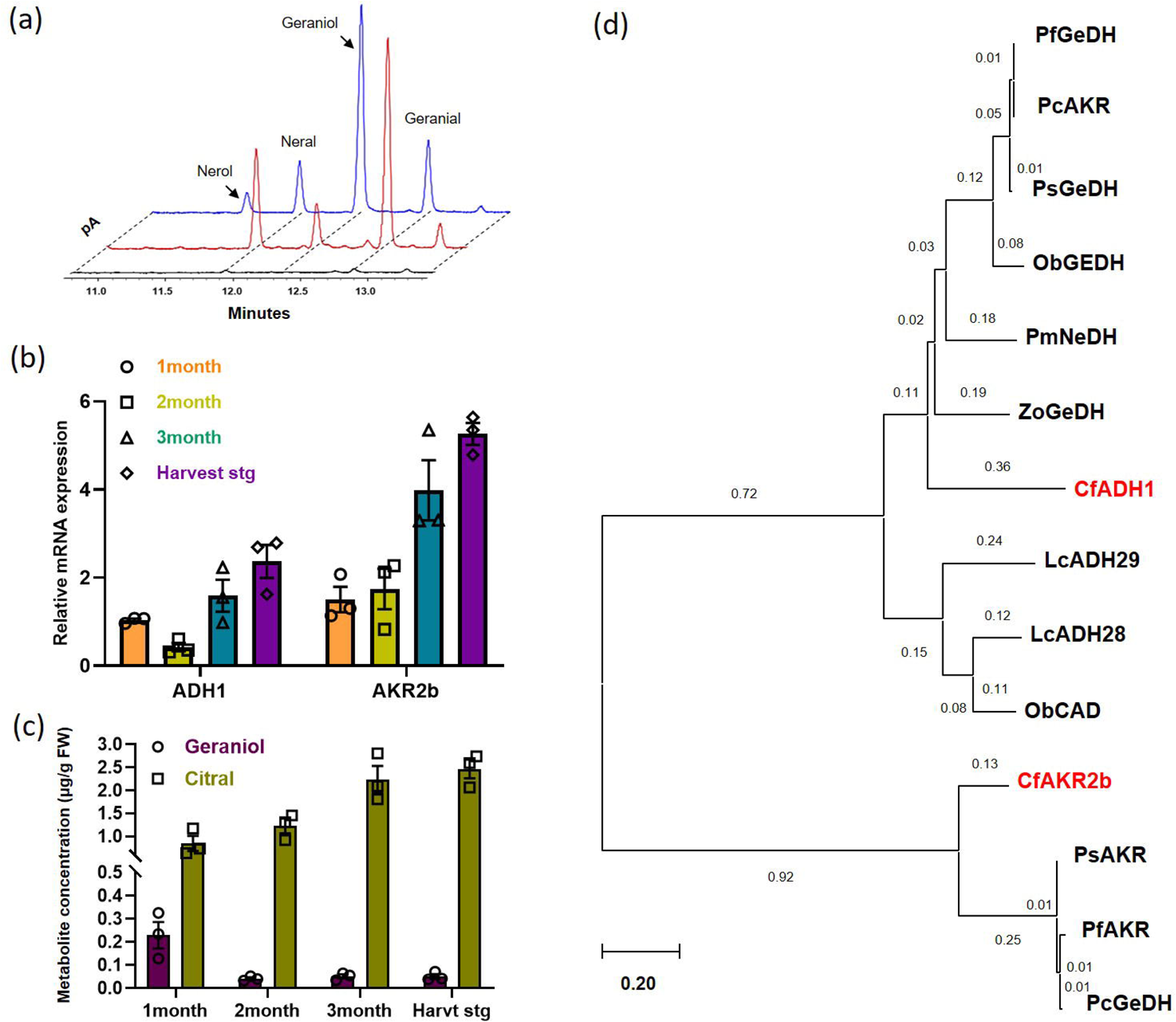
ADH activity in lemongrass leaves, correlation of gene expression and metabolite levels, and phylogentic analyses. (a) Determination of presence of ADH activity in crude total protein extracts oflemongrass leaves. The ADH activity assay was carried out by incubating the leaf crude protein extract in the presence or absence (control) of geraniol as the substrate and NADP cofactor, with a hexane overlay. Reaction products extracted into the hexane fraction were analysed by GC-MS and the products fonned in the assay were compared to the volatile profile of lemongrass leaf. (b) Reverse transcriptase-quantitative polymerase chain reaction (RT-qPCR) analysis showing the relative expression of *CJADHI* and *CJAKR2b* in different developmental stages of lemongrass. Total RNA isolated from leaves was used for RT-qPCR analysis, and the expression level of genes was nonnalized to *CjEFJo.* endogenous control and was set to 1 in CfADHl. (c) Quantification of geraniol and citral content in different developmental stages of lemongrass. Leaves from different developmental stages were harvested and essential oil was distilled and anlyzed by gas chromatography. The amount of geraniol and citral was quantified in relation to the IS. The results shown are from three independent experiments indicated by data points. Statistical analysis was performed by Sidak multiple comparison test using GraphPad Prism 8.0: *\*P<0.05;* **P<0.01; *\*\*\*P<0.001.* Error bars indicate mean± SE. (d) Phylogenetic analysis of CfADHl and CfAKR2b with characterized citral forming enzymes of other plants. Abbreviations: LcADH, *Litsea cubeba ADH,·* CmADH, *Cinnamo111um 111icranthurn ADH,·* PmNeDH, *Persicaria minor nerol dehydrogenase;* ZoGeDH, *Zingiber officinale geraniol dehydrogenase,·* ObCAD, *Ocinnm, basilicum cinnamoyl alcohol dehydrogenase;* ObGEDH, *Ocimum basilicum geraniol dehydrogenase,·* PfAKR, *Perilla frutescens aldoketoreducatse;* PfGeDH, *Perilla frutescens geraniol dehydrogenase,·* PcAKR, *Perilla citriodora aldoketoreducatse;* PcGeDH, *Perilla citriodora geraniol dehydrogenase;* PsAKR, *Perilla setoyensis aldoketoreducatse;* PsGeDH *Perilla setoyensis geraniol dehydrogenase*.

To evaluate the role of *CfADH1* and *CfAKR2a* in citral biosynthesis, a correlative analysis of gene expression and metabolites was performed across various developmental stages (1 month, 2 months, 3 months, and harvesting) of lemongrass leaves. *CfAKR2b* exhibited a gradual increase in expression from the 1^st^ month to the harvesting stage, while *CfADH1* initially decreased in the 2^nd^ month and then increased thereafter. Notably, *CfAKR2b* consistently displayed higher expression levels than *CfADH1* across all stages, with the 3^rd^ month and harvesting stages showing nearly double the expression levels of *CfADH1* (Fig. 1b). Further, determination of geraniol and citral levels in different stages revealed that geraniol content peaked in the 1st month but drastically declined in the subsequent stages, whereas citral content was at its lowest in the 1st month and exhibited a steady and drastic increase in the 2nd, 3rd, and harvesting stages (Fig. 1c). This pattern of gene expression and metabolite levels suggested a potential link between *CfADH1*, *CfAKR2a*, and citral production.

The pattern of gene expression and metabolites correlated with each other indicating their possible role in utilization of geraniol for citral production (Figs 1b, 1c). Further, to determine the evolutionary relatedness of CfADH1 and CfAKR2b, a phylogenetic analysis was performed with other characterized enzymes shown to catalyse the formation of citral from geraniol. CfADH1 was clustered in the group that has other enzymes having specific GeDH activity and exhibited highest amino acid identity of 59% with *Zingiber officinale* (ZoGeDH1) that catalyzes bidirectional inter-conversion of geraniol to citral followed by *Persicaria minor* PmNeDH with 56.58% of amino acid identity (Iijima *et al*., 2014; Tan *et al*., 2018). CfADH1 showed least amino acid identity to *L. cubeba* LcADH29, which was shown to form citral. Whereas CfAKR2b grouped together with *Perilla* AKRs that have been shown to catalyse geraniol to citral conversion under *in vitro* conditions. CfAKR2b exhibited 67-68 % similarity to *Perilla* AKRs (Sato-Masumoto & Ito, 2014).

### Enzymatic characterization of CfADH1 and CfAKR2a

To assess the biochemical nature of CfADH1 and CfAKR2a enzymes, they were expressed in *E. coli* and the purified recombinant proteins were assessed for their enzyme activity. Analysis of enzymatic products by GC/GC-MS analysis showed that CfADH1 catalyzed the oxidation reaction to form citral (geranial and neral) from geraniol with NADP as cofactor (Fig. 2). On the other hand, CfAKR2b exhibited a similar enzymatic activity to that of CfADH1 with a slight difference in terms of products. When incubated with geraniol and NADP cofactor, CfAKR2b formed citral, similar to CfADH1, but also produced an additional product, nerol albeit at a lower level than that of neral and geranial (Fig. 3). As nerol is a *cis*-isomer of geraniol, assays were performed to assess the ability of CfADH1 and CfAKR2b to catalyse the formation of citral from nerol. While CfADH1 did not show any activity with nerol substrate and NADP cofactor, CfAKR2b was able to convert nerol to citral and geraniol in presence of NADP (Fig. S4). As ADHs are known to catalyse redox reactions, assays were performed using citral as the substrate with NADPH as the cofactor, which revealed that CfADH1 could catalyze the reduction of citral into geraniol. Conversely, CfAKR2b could catalyze the reduction of citral into both geraniol and nerol in presence NADPH as the cofactor (Fig S5).

**Figure 2.**
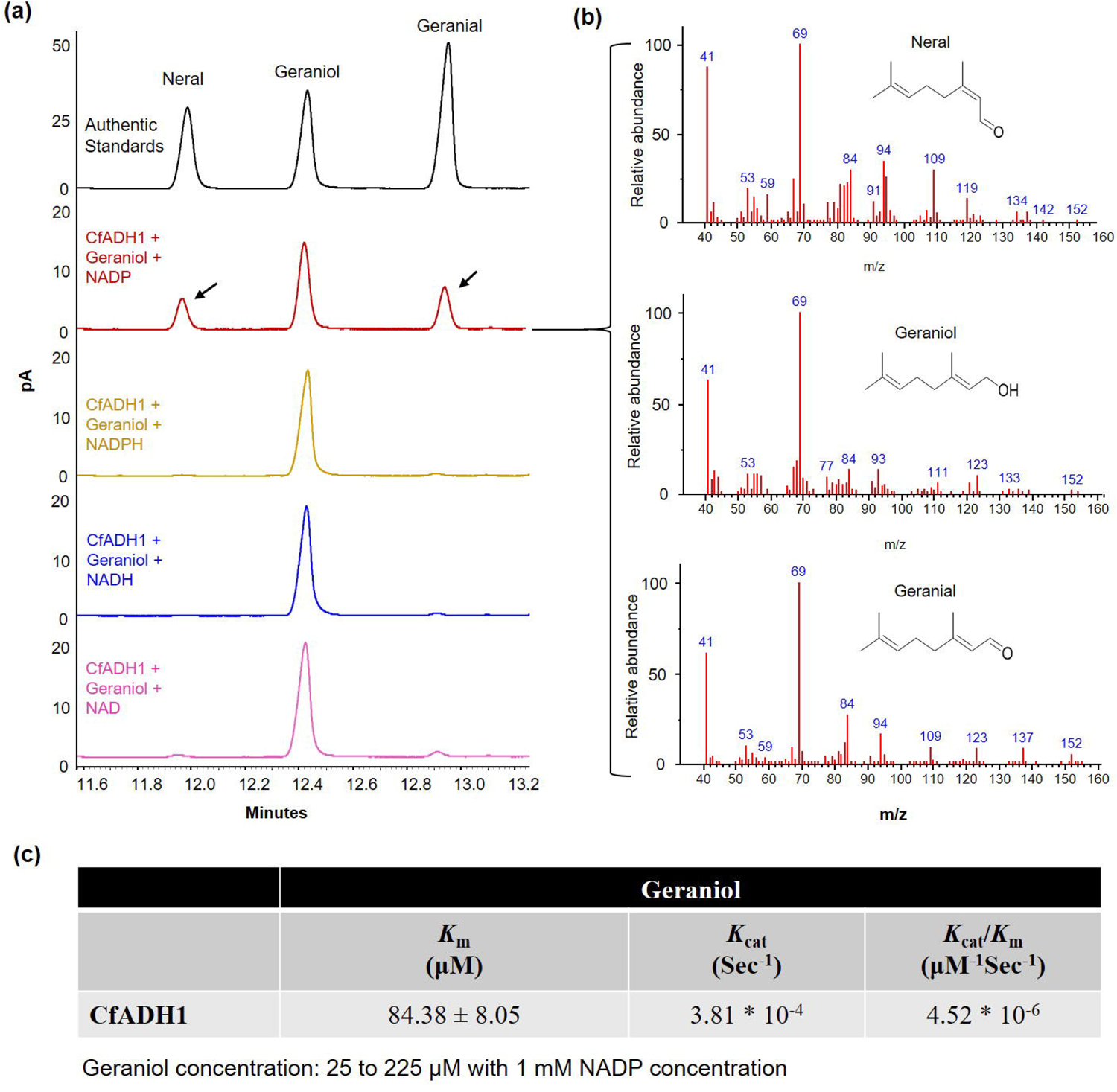
Biochemical characterization of recombinant CfADHl enzyme. (a) Analysis of reaction products formed by recombinant CfADHl. Gas Chromatography-Mass Spectrometry (GC-MS) profile showing the products formed by CfADHl from geraniol. The ADH activity assay was canied out by incubating the purified recombinant CfADHl in the presence of geraniol as the substrate and cofactor, with a hexane overlay. Reaction products extracted into the hexane fraction were analysed by GC-MS and the products fom1ed in the assay were confirmed by comparing the retention time of authentic citral standard. (b) Mass spectra of substrate and product peaks of the assay. The products formed in the assay were further confirmed by comparing the mass spectra of authentic citral standard. (c) Kinetic parametrs of CfADHl. Enzyme kinetics was dete1mined using varying concentrations of geraniol with constant concentration ofNADP.

**Figure 3.**
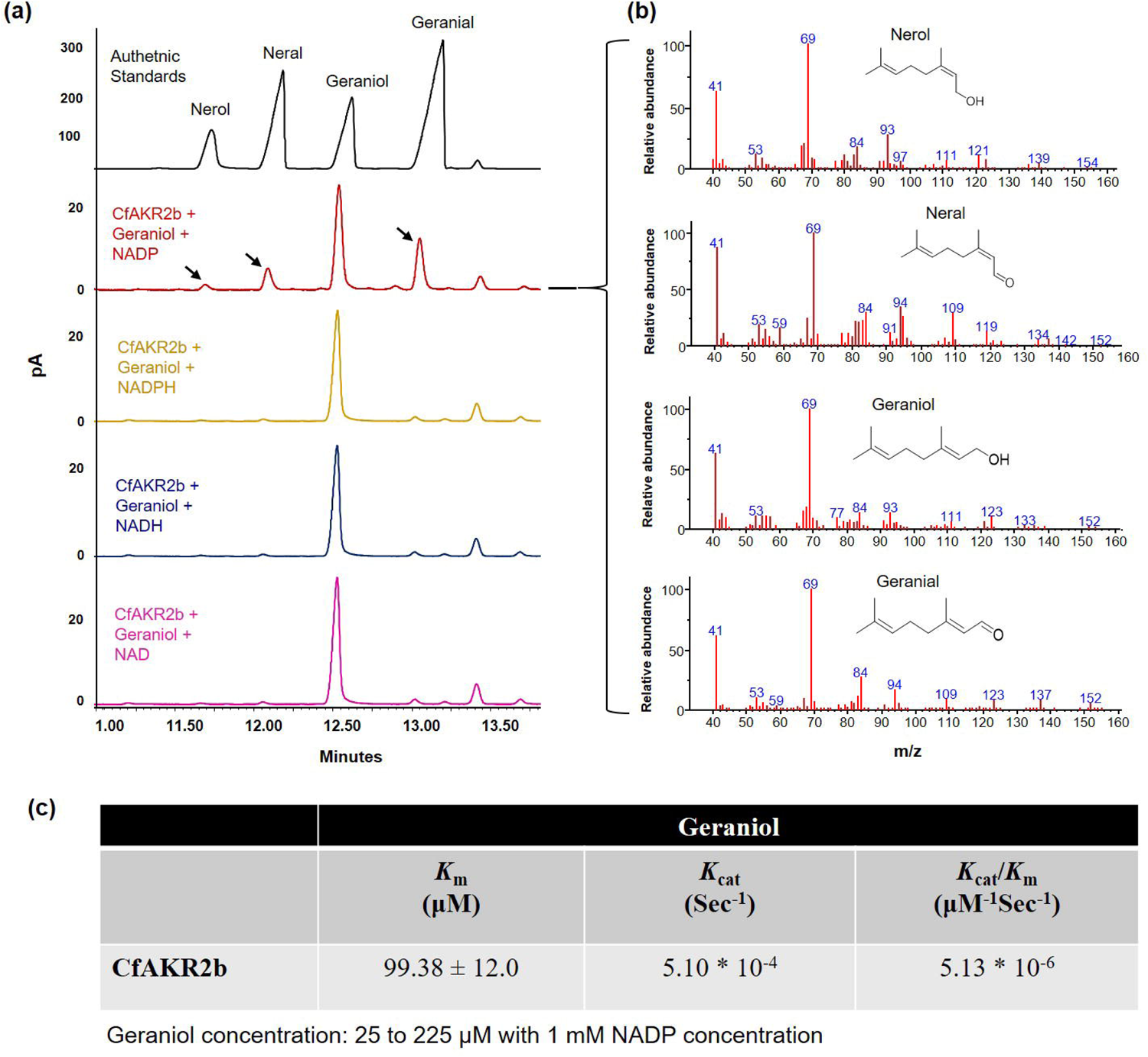
Enzymatic characterization of recombinant CfAKR2b enzyme. (a) Detem1ination of reaction products fonned by recombinant CfAKR2b. Gas Chromatography-Mass Spectrometry (GC-MS) profile showing the products formed by CfAKR2b from geraniol. The enzyme activity assay was can-ied out using the purified recombinant CfAKR2b in the presence of geraniol as the substrate and cofactor, with a hexane overlay. Reaction products fonned in the assay and unreacted substrate were extracted into the hexane fraction and analysed by GC-MS. The products fom1ed in the assay was confirmed by comparing the retention time of authentic citral standard. (b) Mass spectra of substrate and product peaks of the assay. The products formed in the assay were further confirmed by comparing the mass spectra of authentic citral standard. (c) Kinetic parametrs of CfAKR2b. Enzyme kinetics was determined using varying concentrations of geraniol with constant concentration of NADP.

To investigate the enzyme activity of CfADH1 and CfAKR2b with different cofactors, assays were conducted using geraniol or citral as substrates along with different cofactors. Neither CfADH1 nor CfAKR2b displayed any activity when alternative cofactors like NAD or NADH or NADPH were employed with geraniol as the substrate. Likewise, neither enzyme exhibited activity in the reverse reaction using citral as a substrate and NADP or NADH or NAD as cofactors. Additionally, both CfADH1 and CfAKR2b did not show enzyme activity with other alcohol substrates such as linalool, patchoulol, menthol, elemol when using NADP cofactor, confirming their specific geraniol dehydrogenase (GEDH) activity (Fig. S6). Interestingly, despite their sequence un-relatedness, both enzymes exhibited similar *in vitro* activity in terms of geraniol to citral formation with about 40 to 45 % conversion of geraniol to citral with NADP (Figs 2, 3).

Furthermore, it was observed that only CfAKR2b was able to catalyze the conversion of citronellal to citronellol in the presence of NAPDH, whereas there was a trace level of activity when NADH and NADP were used as cofactors (Fig. S7). Further, it was revealed that CfAKR2b could form both *S* and *R* enantiomers of citronellol from their respective citronellal enantiomers (Fig. S8). However, no activity was observed with citronellol as a substrate, along with none of the cofactors used (Fig. S9). The kinetic characterization of CfADH1 and CfAKR2b using varying concentrations of geraniol (25–225 µM) in the presence of constant NADP (100 µM) revealed a similar and comparable pattern between the two enzymes. CfADH1 exhibited an apparent *K*_m_ value of 84.38 µM, whereas CfAKR2b displayed a marginally elevated *K*_m_ of 99.38 µM (Figs 2c, 3c). CfAKR2b possessed a slightly higher catalytic efficiency of 5.13 x 10^-6^ compared to CfADH1, which had 4.52 x 10^-4^.

### VIGS of *CfADH1* and *CfAKR2b* affect citral accumulation in lemongrass

Before taking up silencing of *CfADH1* and *CfAKR2b*, the protocol for virus-induced gene silencing (VIGS) was established in lemongrass. Since TRV-mediated VIGS has been successfully demonstrated in other monocots of Poaceae family such as wheat and maize (Zhang *et al*., 2017; Mei *et al*., 2019), we attempted a similar approach. As PDS is widely used as a marker in VIGS, a 551-bp *PDS* fragment (GenBank accession No. PP421065) from lemongrass leaf cDNA amplified and cloned into *Bam*HI site of pTRV2 vector. The resulting TRV:*CfPDS* construct was mobilized into *Agrobacterium tumefaciens*. Lemongrass seedlings (4 to 5-leaf stage of one month old) were infected with *A. tumefaciens* harbouring TRV:*CfPDS* and empty vector by vacuum-infiltration (Purkayastha *et al*., 2010) as well as through injection of *Agrobacterium* solution into tiller-base region of seedling which facilitated percolation of solution all the way up to leaf. While most of the vacuum-infiltrated seedlings died after few days, most syringe infiltrated seedlings survived and showed the photobleached phenotype in newly emerging leaves 30-40 days postinfiltration (dpi). The phenotype ranged from mild photobleaching with yellowish color to strong photobleaching in the entire leaf (Fig. S10). Subsequent qPCR analysis in corresponding leaves showed a significant decrease in *PDS* mRNA levels in tissues of *CfPDS*-vigs plants when compared with EV control supporting the observed albino phenotype in *CfPDS*-vigs plants (Fig. S10)

Having established a VIGS protocol in lemongrass seedlings, we made sure that the stage of the seedling used for VIGS produces similar metabolite profile to that of mature plants. Next, the same method was adopted to down-regulate the expression of *CfADH1* and *CfAKR2b* in lemongrass leaves to determine their *in planta* role in citral biosynthesis. To prevent potential cross-silencing of *CfADH* and *CfAKR* isoforms, regions of *CfADH1* and *CfAKR2b* that showed the highest dissimilarity with other ADH and AKR isoforms were utilized to generate VIGS constructs (Figs S11, S12). In parallel, the pTRV2:*CfPDS* construct was employed to silence the *PDS* gene, leading to a photobleached phenotype that served as a visible marker to determine the optimal timing for collecting silenced leaf tissues from *CfADH1*-vigs and *CfAKR2b*-vigs plants, ensuring homogeneity in the harvested plant material (Fig. 4a). Moreover, the silencing of *CfADH1* and *CfAKR2b* did not result in any obvious visible phenotype (Fig. 4a). However, typical viral infection phenotype such as slight curling of leaves and patches were observed in infected leaves. Analysis of gene expression by RT-qPCR revealed that the silencing of *CfADH1* and *CfAKR2b* resulted in a significant downregulation of their transcript levels in the range of 56-83% in newly emerging leaves of silenced plants compared to the empty vector (EV) control (Fig. 4b). To evaluate the effect of silencing of these genes on citral accumulation, volatiles were extracted from leaves and analysed. The results from GC analysis showed a significant reduction in the levels of citral (Fig. 4d). The leaves of *CfADH1*-vigs plants produced 37.48% lower levels of citral than that of EV leaves (Fig. 4d). This was followed by a slight increase in the levels of geraniol (54.8%). Similarly, *CfAKR2b*-vigs leaves showed a drastic and significant decline in citral (43.7%), followed by a non-significant increase in geraniol content as compared to EV control leaves (Fig. 4d). Though the degree of *CfADH1* silencing was higher than that of *CfAKR2b* silencing, the level of citral reduction relative to gene expression was greater in the case of *CfAKR2b*-vigs samples (Fig. 4).

**Figure 4.**
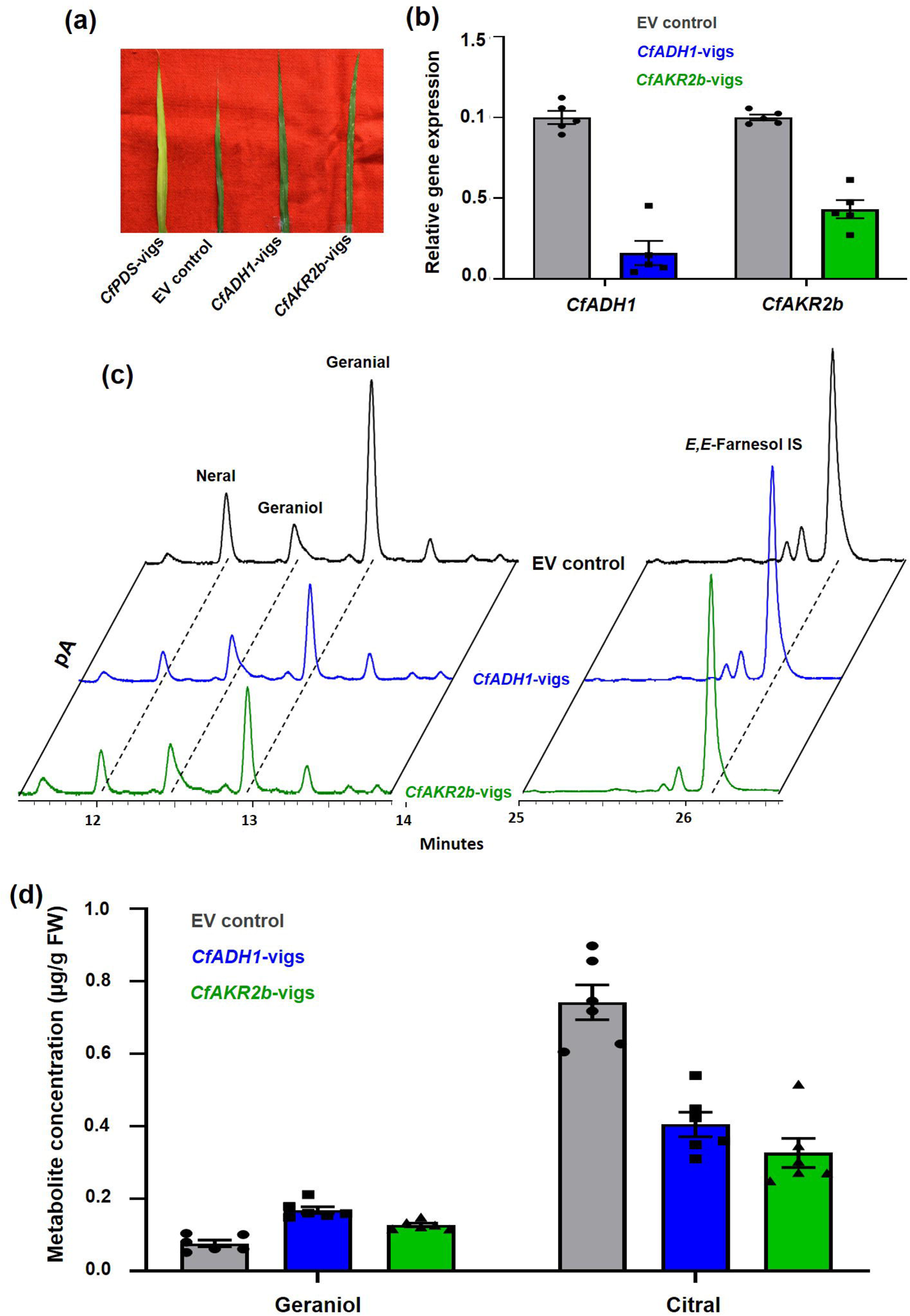
Virus-induced gene silencing (VIGS) of *CfADHJ* and *CfAKR2b* and its effect on citral biosynthesis in lemongrass. (a) Phenotype of leaves infected with empty vectors (EV control), pTRV2::CfPDS, *pTRV2::CfADH1,* and *pTRV2::CfAKR2b.* (b) Reverse transcriptase-quantitative polymerase chain reaction (RT-qPCR) analysis showing the relative expression of *CfADHl,* and *CfAKR2b* in respective VIGS samples. Total RNA isolated from EV control, *CfADHJ-vigs,* and *CfAKR2b-vigs* leaves was used for RT-qPCR analysis. The expression level of genes was normalized to *CfEF1a* endogenous control and was set to 1 in EV control to determine the relative reduction of transcripts in *CfADHl-vigs,* and *CfAKR2b-vigs* leaves. (c) Representative chromatograms from EV control, *CfADHl-vigs,* and *CfAKR2b-vigs* showing geraniol and citral peaks. Volatiles were extracted using hexane containing internal standard (IS) E,E-famesol, and subjected to gas chromatography (GC) analysis. (d) Quantification of geraniol and citral in EV control, *CfADHl-vigs,* and *CfAKR2b-vigs* tissues. The amount of geraniol and citral was quantified in relation to the IS. The results shown are from five to six independent experiments indicated by data points. Statistical analysis was perfonned by Sidak multiple comparison test using GraphPad Prism 8.0: *\*P<0.05; **P<0.0l; ***P<0.001.* Error bars indicate mean ± SE.

### Expression of *CfADH1* and *CfAKR2b* increases citral production in lemon balm

Suppression of *CfADH1 and CfAKR2b,* negatively affected the citral biosynthesis in lemongrass (Fig 4). To further determine the ability of CfADH1 and CfAKR2b to form citral in leaf physiological conditions, the coding sequence of these genes was placed under the control of the cauliflower mosaic virus (CaMV) 35S promoter, resulting in overexpression constructs. Since the development of transgenic lemongrass is not feasible due to non-availability of efficient regeneration and transformation method, and moreover, agroinfiltration in lemongrass leaves is not possible due to their thick and hygroscopic nature, lemon balm leaves that produce a similar monoterpene profile to that of lemongrass were used for transient expression by agroinfiltration (Fig. 5a). Analysis of gene expression 96 h after agroinfiltration showed that while pRI::*CfADH1* infiltrated lemon balm leaf samples had an 2.4-fold increase in *CfADH1* expression, those of pRI::*CfAHR2b* samples had a 3-fold expression of *CfAHR2b* (Fig. 5b). Subsequent analysis of metabolites by GC analysis revealed an elevation in the levels of citral in both *CfADH1* and *CfAKR2b* expressing leaves compared with the EV control as observed in the chromatogram (Fig. 5c). Further quantification of metabolites clearly showed a significant increase in citral level in both *CfADH1* and *CfAKR2b* expressing lemon balm leaves. While *CfADH1* overexpressing samples showed 1.9 fold fold increased accumulation of citral, *CfAKR2b* overexpression led to 2.19 fold increased citral production (Fig. 5d). Besides, both *CfADH1* and *CfAKR2b* expressing leaves showed a slight decline in the level of geraniol (Fig. 5d).

**Figure 5.**
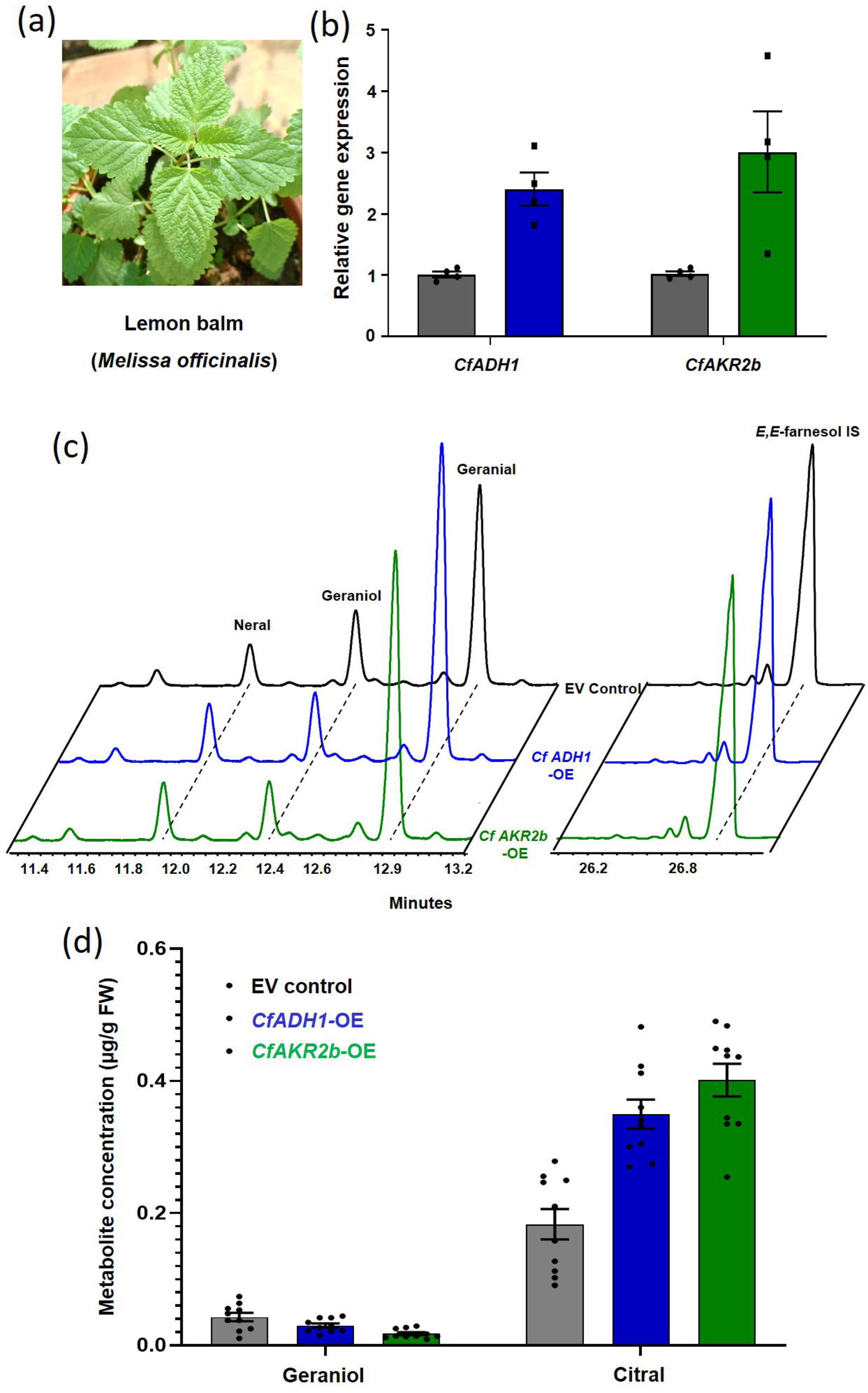
Transient expression of *CfADHJ* and *CfAKR2b* and its effect on citral biosynthesis in lemon balm *(Melissa officinalis).* (a) Representative image of lemon balm plant used for transient expression. (b) Reverse transcriptase-quantitative polymerase chain reaction (RT­ qPCR) analysis showing the relative expression of *CfADHJ* and *CfAKR2b* in respective overexpression samples. Total RNA isolated from empty vector (EV control), *CfADHJ-OE* and *CfAKR2b-OE* leaves was used for RT-qPCR analysis. The expression level of genes was nonnalized to *CfEFla* endogenous control and was set to l in EV control to detennine the relative level of transcripts in *CfADHJ-OE* and *CfAKR2b-OE* leaves. (c) Representative chromatograms from EV control, *CfADH1-OE,* and *CfAKR2b-OE* showing geraniol and citral peaks. Volatiles were extracted using hexane containing internal standard (IS) E,E-famesol, and subjected to gas chromatography (GC) analysis. (d) Detennination of geraniol and citral content in lemon balm leaf tissues infiltrated with *Agrobacterium* harbouring EV control, *CfADHJ-OE,* and *CfAKR2b-OE* constructs. The amount of geraniol and citral was quantified in relation to the IS. The results shown are from four (RT-qPCR) to ten (metabolites) independent experiments indicated by data points. Statistical analysis was perfon11ed by Sidak multiple comparison test using GraphPad Prism 8.0: \**P<0.05;* **P<0.01; *\*\*\*P<0.001.* EITor bars indicate mean± SE.

### Subcellular localization of CfADH1 and CfAKR2b

Geraniol, the substrate for citral is known to be biosynthesized in mostly plastids by the action of geraniol synthase, but recent studies in some plants such as rose and geranium show that it is formed in cytoplasm by NUDX hydrolases. Monoterpenes are generally made in plastids, however, recent studies indicate in some species they are formed through cytosolic pathway (Gutensohn *et al*., 2013; Bergman *et al*., 2021a,b; Conart *et al*., 2022). Hence, to determine the subcellular location of CfADH1 and CfAKR2b, the enzymes responsible for citral biosynthesis in lemongrass, an *in silico* analysis was performed utilizing six different prediction software programs.

In the case of CADH1, three programs indicated plastidial localization, two programs predicted cytosolic localization, and one program indicated a dual plastidial and cytosolic localization. Whereas for CfAKR2b, four out of the six programs predicted cytosolic localization, while two programs predicted plastidial location (Table S3). As *in silico* prediction programs did not yield a consistent localization pattern for CfADH1 and CfAKR2b, we assessed their subcellular localization experimentally by agroinfiltrating yellow fluorescent protein (YFP) fusion constructs into *N. benthamiana* leaves. Analysis of fluorescence in infiltrated leaves by confocal microscopy revealed a diffusing signal in the cell, akin to the fluorescence observed in cytosolic YFP control, indicating cytosolic localization for CfADH1-YFP infiltrated samples. Intriguingly, fluorescence was also observed as certain dots, suggesting plastidial localization (Fig. 6). Moreover, the overlapping signals emitted by the chlorophyll autofluorescence (AF) and the YFP signal from CfADH1-YFP provided clear evidence that CfADH1 is dually localized to the cytoplasm and plastids (Figs 6, S13). In the case of CfAKR2b-YFP, the fluorescence was exclusively associated with the cytoplasm, resembling the fluorescence from the YFP control sample. Unlike CfADH1-YFP, there was no overlap of any signal with chlorophyll autofluorescence confirming the cytosolic nature of CfAKR2b (Fig. 6).

**Figure 6.**
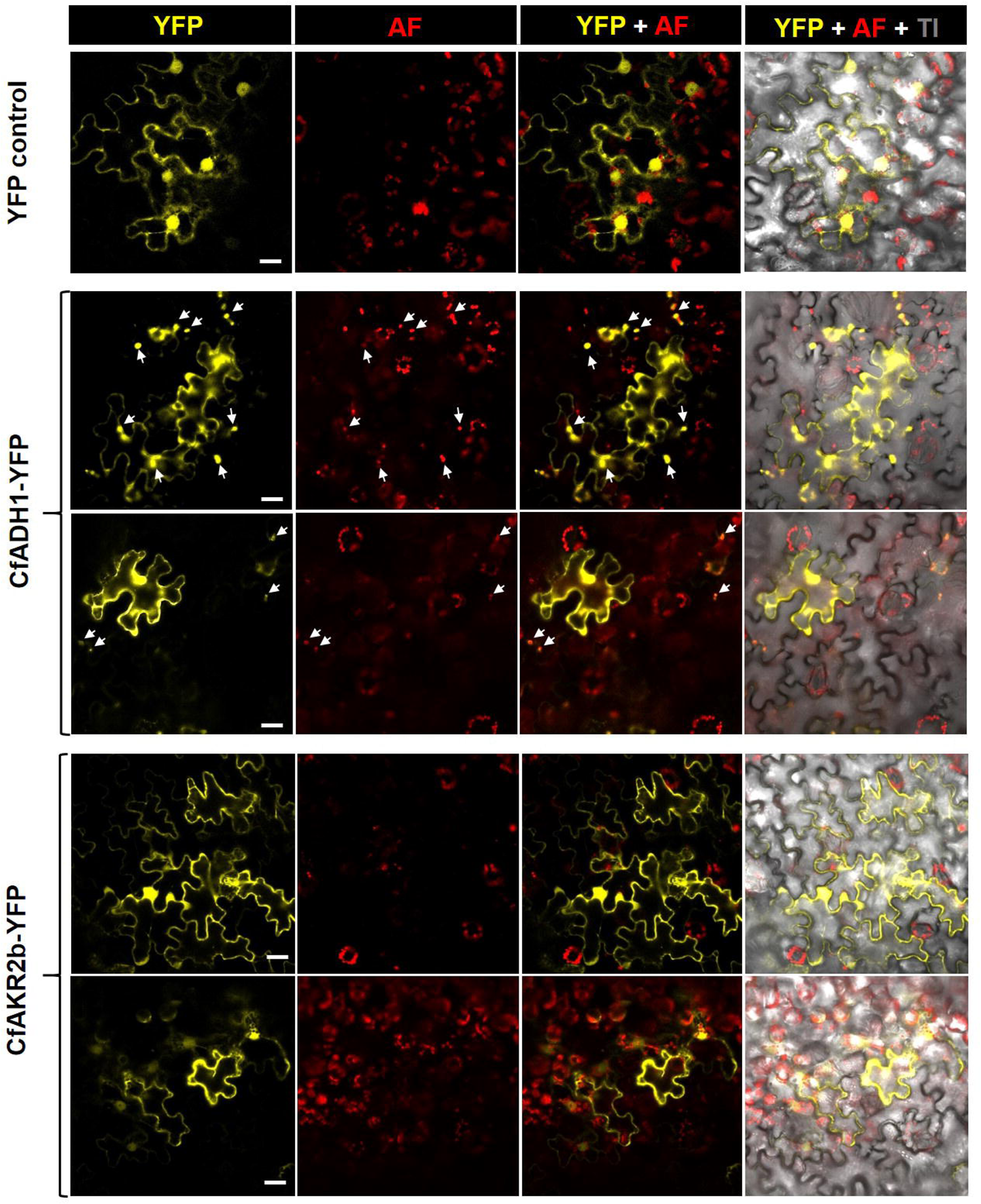
Analysis of subcellular localization of CfADH1 and CfAKR2b. Confocal laser scanning microscopy of *Nicotiana benthamiana* leaves transformed with CfADHl-YFP and CfAKR2b-YFP expression constructs. YFP fluorescence and chlorophyll autoflourescence are shown in YFP and AF columns, respectively. The third vertical panel shows merged image of YFP and AF, and the fourth and the rightmost panel shows merged YFP, AF, and transmission image. The arrows indicate chloroplasts. Scale bar: 20 µm.

### Effect of fosmidomycin and lovastatin on geraniol and citral production

Our subcellular localization along with *in planta* findings strongly supported the notion that citral biosynthesis involves the concerted action of dual-localized enzymes, CfADH1 in the plastid-cytosol and CfAKR2b in the cytosol, indicating a dual origin of citral from both cellular compartments. To further corroborate this, lemongrass seedlings were treated with lovastatin (1 mM), a potent inhibitor of the MVA pathway, and fosmidomycin (2 mM), a specific inhibitor of the MEP pathway, and monitored volatile compound production after 48 hours. Both inhibitors led to significant reductions in geraniol and citral production (Fig. 7). Interestingly, lovastatin treatment resulted in a slightly greater reduction in geraniol and citral levels (60-62%) compared to fosmidomycin treatment, which affected approximately 53-55% of geraniol and citral levels (Fig. 7). These findings further supported the subcellular localization results and provided evidence for the dual cytosolic- and plastidial-origin of citral biosynthesis. The results showed that citral is formed via both the cytosolic mevalonic acid (MVA) and plastidial methylerythritol phosphate (MEP) pathways in lemongrass.

**Figure 7.**
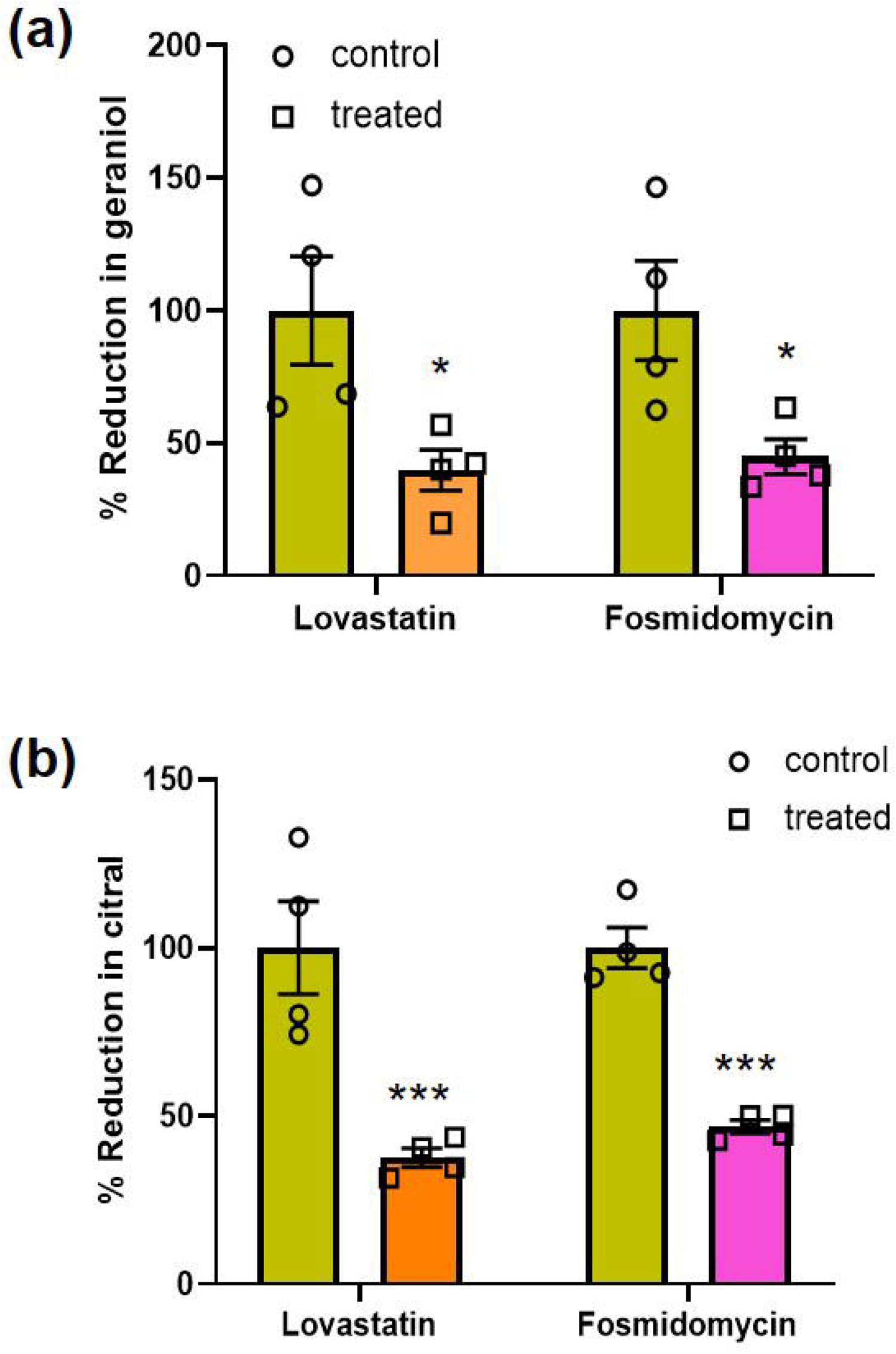
Effect of lovastatin and fosmidomycin feeding on geraniol and citral accumulation. Young 4-leaf stage seedlings were cut at the tiller base region and were incubated in 500 µL of an aqueous solution of fosmidomycin (2 mM) or 500 µL of a 20% ethanolic solution of lovastatin (1 mM) with appropriate controls. After 48 h of incubation in 16h light and 8 h dark condition, the volatiles were extracted using hexane containing the internal standard *(E,E­* farnesol) and analysed by GC analysis. The results shown are from four independent samples. Statistical analysis was performed by Sidak multiple comparison test using GraphPad Prism 8.0: \**P<0.05;* ***P<0.001. Error bars indicate mean± SE.

## Discussion

In plants, it was widely accepted that monoterpenes are synthesized within plastids via the methylerythritol phosphate (MEP) pathway, catalyzed by monoterpene synthases. However, this understanding has been challenged in a few plants where monoterpene synthases have been found in the cytosol (Aharoni *et al*., 2004; Dong *et al*., 2016). Additionally, bifunctional cytosolic mono-/sesquiterpene synthases have been identified, capable of producing monoterpenes (Aharoni *et al*., 2004; Nagegowda *et al*., 2008; Davidovich-Rikanati *et al*., 2008). Further, in recent years an unexpected enzyme belonging to a NUDX hydrolase was shown to be involved in formation of monoterpene geraniol in the cytosol (Magnard *et al*., 2015; Bergman *et al*., 2021a; Zhou *et al*., 2022). These studies indicated the contrasting evolution of monoterpene biosynthesis in different plant species. Citral, the monoterpene aldehyde found as major component of essential oils of many plants, is derived mostly from geraniol or in some cases nerol by alcohol dehydrogenases (ADHs). While enzymes of the ADH class in few plant species have been biochemically characterized, showcasing their ability to catalyze the conversion of geraniol and/or nerol into citral under *in vitro* conditions (Iijima *et al*., 2006, 2014; Lüddeke *et al*., 2012; Sato-Masumoto & Ito, 2014a; Tan *et al*., 2018), there exists only one instance providing both biochemical and genetic evidence for the participation of ADHs in citral formation within *L. cubeba* (Zhao *et al*., 2024). Despite these studies, there is still limited knowledge on citral biosynthesis and its origin in plants, and especially in lemongrass, a citral-rich essential oil producing monocot species. Here, we provide biochemical and genetic evidences that two phylogenetically distant and differentially localized CfADH1 and CfAKR2b are involved in citral biosynthesis in lemongrass.

The expression of both *CfADH1* and *CfAKR2b* exhibited a positive correlation with the level of citral in different developmental stages, indicating their involvement in citral biogenesis. CfADH1 exhibited highest amino acid identity with ZoGeDH followed by and PmNeDH from among the characterized ADHs, while CfAKR2b showed homology with AKRs from *Perilla* (Fig. 1d) (Iijima *et al*., 2014; Sato-Masumoto & Ito, 2014; Tan *et al*., 2018) Both CfADH1 and CfAKR2b possessed similar enzyme abilities to catalyse the formation of citral in the presence of NADP. This ability of CfADH1 and CfAKR2b was similar to the enzyme activity found in the crude leaf protein extract, and the product (citral) profile of these matched the volatile profile of lemongrass leaf (Fig. 1a). While geranial formation was always higher than the formation of neral (about 2:1 ratio), which corresponds to the ratio of geranial:neral found in volatile profile of lemongrass. This property of CfADH1 and CfAKR2b is similar to the enzymatic ability of ADHs and AKRs from *Perilla* spp. (Sato-Masumoto & Ito, 2014). The kinetic parameters of both enzymes were similar, with CfAKR2b having slightly higher *K*_m_ and *K*_cat_ than that of CfADH1 (Figs 2, 3). Also, the *K*_m_ of both enzymes were in a similar range to that of previously characterized monocot ZoGeDH (*K*_m_ = 60.9) from ginger that catalysed the conversion of geraniol to citral (Iijima *et al*., 2014). Interestingly, CfAKR2b also exhibited the ability to reduce citronellal to its alcohol citronellol, which was also observed in the assay with crude protein extract of lemongrass leaves (Fig. S7). However, the volatile profile of lemongrass leaves did not have the presence of either citronellal or citronellol, indicating that this ability could be due to the *in vitro* promiscuous activity and may not be an *in vivo* scenario.

To further validate the *in planta* role of CfADH1 and CfAKR2b in lemongrass citral biogenesis, we established an *in planta* gene assay by VIGS. TRV-based silencing has been achieved widely in dicot species, but there are only few reports of its use in monocots, where it has been successfully utilized in rice, maize and wheat to study the gene function (Purkayastha *et al*., 2010; Kirigia *et al*., 2014; Kant *et al*., 2015; Zhang *et al*., 2017). Since TRV has a broad host range and could infect maize and wheat, we predicted that it could also infect lemongrass as it is from the same family, and accordingly different infection procedures were attempted. Among the different methods tried, we got success in silencing *PDS* gene by agroinjection method (Fig. S10), which was subsequently used for silencing our target genes i.e., *CfADH1* and *CfAKR2b*. The silencing of either *CfADH1* or *CfAKR2b* resulted in a significant reduction in citral content, accompanied by a slight increase in its substrate, geraniol (Fig. 4). Interestingly, despite *CfAKR2b* being silenced to a lesser extent (56.8%) compared to *CfADH1* (83%), there was a greater impact on citral accumulation. This may be attributed to the higher efficiency of CfAKR2b, which possesses a greater enzyme turnover number than CfADH1 (Fig. 3c). To substantiate the results observed in VIGS, we overexpressed *CfADH1* and *CfAKR2b* transiently in lemon balm, a dicot plant known to produce a citral-rich volatile profile similar to that of lemongrass, with citral contributing to over 80% of its composition (Kittler *et al*., 2018). We used lemon balm as lemongrass is not amenable to both transient and stable transformation using *Agrobacterium*. The transient expression of *CfADH1* and *CfAKR2b* resulted in a significant increase in the accumulation of citral, accompanied by a notable decline in the levels of geraniol in lemon balm (Fig. 5). These results not only reaffirmed the *in planta* involvement of CfADH1 and CfAKR2b enzymes in citral production but also indicated the possibility of a similar scenario of citral biosynthesis in lemon balm to that observed in lemongrass. Indeed, our assembly of the lemon balm transcriptome (Mansouri & Mohammadi, 2021) and subsequent blast search revealed the presence of gene candidates encoding proteins with similarity to CfADH1 and CfAKR2b (Fig. S9). Moreover, the assembly of publicly available transcriptome data sets from other citral-producing plant species such as *Verbena officinalis* (Li et al., 2023) and *L. cubeba* (Zhao *et al*., 2024), and subsequent blast analysis revealed the presence of candidates showing a high degree of similarity to CfADH1 and CfAKR2b with high *in silico* expression values (Fig. S14). This suggests that similar to lemongrass, plants producing citral-rich essential oil may employ a shared strategy of engaging phylogenetically distant ADH and AKR enzymes for citral biosynthesis. However, it is worth noting that though two ADHs (LcADH28 and LCADH29) have been functionally characterized to be involved in citral formation in *L. cubeba* (Zhao et al., 2024), the involvement of AKR-type enzyme identified from the transcriptome assembly cannot be ruled out and needs further investigation. Furthermore, the biochemical characterization of ADH and AKR enzymes from *Perilla*, capable of catalyzing the conversion of geraniol into citral, supported their potential role in citral biosynthesis. However, the *in planta* function of these enzymes was never investigated (Sato-Masumoto & Ito, 2014).

Monoterpene biosynthesis has been generally considered to occur in plastids, although there have been notable exceptions to this scenario (Ohara, 2003; Aharoni *et al*., 2004; Wu *et al*., 2006; Davidovich-Rikanati *et al*., 2008; Dong *et al*., 2013, 2016). Moreover, the precursor for citral biosynthesis, geraniol, is reported to form through the involvement of either TPS encoding GES or through a noncanonical pathway involving the participation of Nudx enzymes (Bergman *et al*., 2021; Conart *et al*., 2022). This prompted us to investigate the subcellular localization of CfADH1 and CfAKR2b. Interestingly, CfADH1 was found to be dual-localized in both the cytosol and plastids, while CfAKR2b was exclusively cytosolic (Fig. 6). In plant terpenoid biosynthesis, enzyme isoforms with various subcellular localizations have been documented for FPPS, IDI, and certain TPSs (Nagegowda and Gupta, 2020). However, instances of dual targeting of the enzyme encoded by the same gene are scarce and have only been reported in the case of nerolidol/linalool synthase from sandalwood, which was primarily targeted to plastids, with some degree of localization also observed in the cytosol (Zhang *et al*., 2021). The dual cytosol/plastid localization of CfADH1 and cytosolic localization of CfAKR2b indicated that citral in lemongrass is made in both cytosol and plastids. Therefore, the substrate geraniol required for citral formation in lemongrass could be synthesized in both compartments to be readily available for utilization by CfADH1 and CfAKR2b. This in turn might be accomplished through the participation of cytosolic NUDX, as demonstrated in Rose and geranium, or by cytosolic GES, as evidenced in *Epiphyllum oxypetalum*, or by plastidial GESs (Zhang et al., 2023; Simkin *et al*., 2013). Supporting this above scenario, analysis of lemongrass-specific transcriptomes showed the presence of both TPS (potentially encoding a GES) and NUDX candidates, which could play a role in the formation of geraniol to be made available for CfADH1 in both the cytosol and plastids, as well as for CfAKR2b in the cytosol (Meena *et al*., 2016; Bhat *et al*., 2023). In fact, feeding lemongrass seedlings with the MVA-pathway specific inhibitor lovastatin and the MEP-pathway specific inhibitor fosmidomycin, showed a significant reduction in both geraniol and citral accumulation, providing additional evidence for the crucial roles of both cytosolic MVA and plastidial MEP pathways in citral biosynthesis (Fig. 7. This is consistent with earlier studies involving [2-^14^C]-MVA feeding in *Cymbopogon winterianus* leaves and [1-^13^C]-glucose and fosmidomycin feeding in lemongrass leaves, which suggested the involvement of the MVA pathway and MEP pathway in geraniol and citral formation, respectively (Akhila, 1985; Gupta & Ganjewala, 2015). These results indicate that lemongrass has evolved the mechanism to recruit ADH and AKR type enzymes to efficiently utilize the precursor available in both the compartments generated via both the MVA and MEP pathways, in this case, geraniol, which could be made by the plastidial GES and the cytosolic NUDX (Fig. 8).

**Figure 8.**
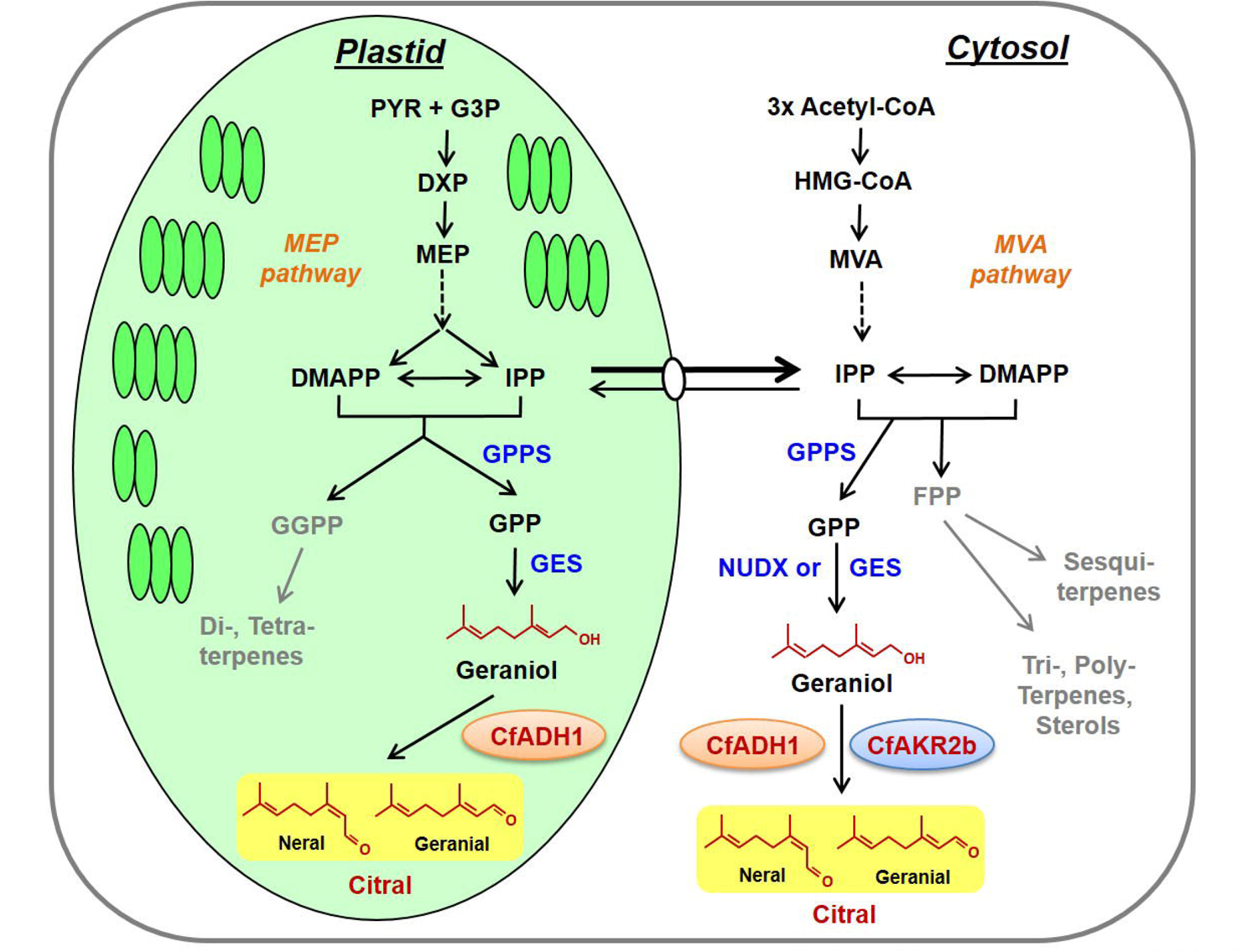
Overview of citral biosynthesis in lemongrass. The dual cytosol/plastid localized CfADHl and cytosolic CfAKR2b fo1m citral by utilizing geraniol made either by plastidial geraniol synthase (GES) or cytosolic NUDX or GES. Abbreviations: DMAPP, dimethylally diphosphate; DXP, 1-Deoxy-D-xylulose 5-phosphate; FPP, farnesyl diphosphate; G3P, glyceraldehyde 3-phosphate; GES; geraniol synthase; GPP, geranyl diphosphate; GPPS, GPP synthase; GGPP, geranylgeranyl diphosphate; HMG-CoA, 3-hydroxy-3-methylglutaryl coenzyme A; IPP, isopentenyl diphosphate; MEP, methylerythritol phosphate; MVA, mevalonic acid; PYR, pyruvate.

With advancements in research, new, non-canonical pathways of terpenoid biosynthesis are being uncovered, expanding our comprehension of the remarkable diversity and complexity within plant terpenoid biochemistry. Extending this knowledge, here we provide biochemical and genetic proof that citral biosynthesis in lemongrass takes place via cytosol and plastids through the involvement of phylogenetically distant dual cytosol/plastid localized ADH and cytosolic AKR enzymes. This study unveils a previously undiscovered mechanism wherein the same terpenoid molecule (i.e., citral) is synthesized by phylogenetically distant enzymes through both the MVA and MEP pathways in lemongrass. This implies an evolutionary strategy aimed at maximizing the utilization of precursor pools from both cytosolic and plastidial pathways, ultimately enhancing citral production.

## Acknowledgements

This research was supported by the MLP0003 and HCP0007 projects of Council of Scientific and Industrial Research. P.G., and A.S. are the recipients of Department of Biotechnology (DBT) and University Grants Commission (UGC) senior research fellowship, respectively. N.R.K was a project associate in MLP0003 and HCP0007 projects. The authors are thankful to Dr. V.S. Pragadheesh for chiral GC analysis and for sparing purified volatiles. Auhtors are also thankful to Drs. Sumit Ghosh for sparing pGWB441 vector, and Rajendra Patel for help with confocal analysis. The authors also express their sincere gratitude to the Director, CSIR-CIMAP for support throughout the study. Institutional communication number for this article is CIMAP/PUB/2024/34.

## Author contributions

DAN planned and designed the research. PG, AS, KNR, PRTK, and RK performed experiments, conducted fieldwork, and analysed data. DAN and PG analysed and interpreted the data and wrote the manuscript.

## Data availability

The authors confirm that the data supporting the findings of this study are available within article in Supporting Information.

## References

Aharoni A, Giri AP, Verstappen FWA, Bertea CM, Sevenier R, Sun Z, Jongsma MA, Schwab W, Bouwmeester HJ. 2004. Gain and Loss of Fruit Flavor Compounds Produced by Wild and Cultivated Strawberry Species. The Plant Cell 16: 3110–3131.

Akhila A. 1985. Biosynthetic relationship of citral-trans and citral-cis in cymbopogon flexuosus (lemongrass). Phytochemistry 24: 2585–2587.

Bergman ME, Bhardwaj M, Phillips MA. 2021a. Cytosolic geraniol and citronellol biosynthesis require a Nudix hydrolase in rose-scented geranium (Pelargonium graveolens). Plant Journal 107: 493–510.

Bergman ME, Bhardwaj M, Phillips MA. 2021b. Cytosolic geraniol and citronellol biosynthesis require a Nudix hydrolase in rose-scented geranium (Pelargonium graveolens). The Plant Journal 107: 493–510.

Bhat S, Sharma A, Sharma P, Singh K, Kundan M, Fayaz M, Wajid MA, Gairola S, Misra P. 2023. Development and analysis of de novo transcriptome assemblies of multiple genotypes of Cymbopogon spp. reveal candidate genes involved in the biosynthesis of aromatic monoterpenes. International Journal of Biological Macromolecules 253: 127508.

Bradford MM. 1976. A Rapid and Sensitive Method for the Quantitation of Microgram Quantities of Protein Utilizing the Principle of Protein-Dye Binding. ANALYTICAL BIOCHEMISTRY 72: 248–254.

Conart C, Saclier N, Foucher F, Goubert C, Rius-Bony A, Paramita SN, Moja S, Thouroude T, Douady C, Sun P, et al. 2022. Duplication and Specialization of NUDX1 in Rosaceae Led to Geraniol Production in Rose Petals (M Purugganan, Ed.). Molecular Biology and Evolution 39.

Davidovich-Rikanati R, Lewinsohn E, Bar E, Iijima Y, Pichersky E, Sitrit Y. 2008. Overexpression of the lemon basil α-zingiberene synthase gene increases both mono- and sesquiterpene contents in tomato fruit. The Plant Journal 56: 228–238.

Deepa K, Sheeja TE, Santhi R, Sasikumar B, Cyriac A, Deepesh P V., Prasath D. 2014. A simple and efficient protocol for isolation of high quality functional RNA from different tissues of turmeric (Curcuma longa L.). Physiology and Molecular Biology of Plants 20: 263–271.

Dong L, Jongedijk E, Bouwmeester H, Van Der Krol A. 2016. Monoterpene biosynthesis potential of plant subcellular compartments. New Phytologist 209: 679–690.

Dong L, Miettinen K, Goedbloed M, Verstappen FWA, Voster A, Jongsma MA, Memelink J, Krol S van der, Bouwmeester HJ. 2013. Characterization of two geraniol synthases from Valeriana officinalis and Lippia dulcis: Similar activity but difference in subcellular localization. Metabolic Engineering 20: 198–211.

Gupta AK, Ganjewala D. 2015. A study on biosynthesis of “citral” in lemongrass (C. flexuosus) cv. Suvarna. Acta Physiologiae Plantarum 37: 240.

Gutensohn M, Orlova I, Nguyen TTH, Davidovich-Rikanati R, Ferruzzi MG, Sitrit Y, Lewinsohn E, Pichersky E, Dudareva N. 2013. Cytosolic monoterpene biosynthesis is supported by plastid-generated geranyl diphosphate substrate in transgenic tomato fruits. The Plant Journal 75: 351–363.

Iijima Y, Koeduka T, Suzuki H, Kubota K. 2014. Biosynthesis of geranial, a potent aroma compound in ginger rhizome (*Zingiber officinale*): Molecular cloning and characterization of geraniol dehydrogenase. Plant Biotechnology 31: 525–534.

Iijima Y, Wang G, Fridman E, Pichersky E. 2006. Analysis of the enzymatic formation of citral in the glands of sweet basil. Archives of Biochemistry and Biophysics 448: 141–149.

Kant R, Sharma S, Dasgupta I. 2015. Virus-Induced Gene Silencing (VIGS) for Functional Genomics in Rice Using Rice tungro bacilliform virus (RTBV) as a Vector. In: 201–217.

Kirigia D, Runo S, Alakonya A. 2014. A virus-induced gene silencing (VIGS) system for functional genomics in the parasitic plant Striga hermonthica. Plant Methods 10: 16.

Kittler J, Krüger H, Ulrich D, Zeiger B, Schütze W, Böttcher C, Krähmer A, Gudi G, Kästner U, Heuberger H, et al. 2018. Content and composition of essential oil and content of rosmarinic acid in lemon balm and balm genotypes (Melissa officinalis). Genetic Resources and Crop Evolution 65: 1517–1527.

Kumar K, Malhotra J, Kumar S, Sood V, Singh D, Sharma M, Joshi R. 2023. Citral enrichment in Lemongrass (Cymbopogon flexuosus) oil using spinning band equipped high vacuum distillation column and sensory evaluation of fractions. Food Chemistry Advances 2: 100291.

Lüddeke F, Wülfing A, Timke M, Germer F, Weber J, Dikfidan A, Rahnfeld T, Linder D, Meyerdierks A, Harder J. 2012. Geraniol and Geranial Dehydrogenases Induced in Anaerobic Monoterpene Degradation by Castellaniella defragrans. Applied and Environmental Microbiology 78: 2128–2136.

Magnard J-L, Roccia A, Caissard J-C, Vergne P, Sun P, Hecquet R, Dubois A, Hibrand-Saint Oyant L, Jullien F, Nicolè F, et al. 2015. Biosynthesis of monoterpene scent compounds in roses. Science 349: 81–83.

Meena S, Kumar SR, Venkata Rao DK, Dwivedi V, Shilpashree HB, Rastogi S, Shasany AK, Nagegowda DA. 2016. De Novo Sequencing and Analysis of Lemongrass Transcriptome Provide First Insights into the Essential Oil Biosynthesis of Aromatic Grasses. Frontiers in Plant Science 7.

Mei Y, Liu G, Zhang C, Hill JH, Whitham SA. 2019. A sugarcane mosaic virus vector for gene expression in maize. Plant Direct 3: 1–13.

Mukarram M, Choudhary S, Khan MA, Poltronieri P, Khan MMA, Ali J, Kurjak D, Shahid M. 2021. Lemongrass Essential Oil Components with Antimicrobial and Anticancer Activities. Antioxidants 11: 20.

Nagegowda DA. 2010. Plant volatile terpenoid metabolism: Biosynthetic genes, transcriptional regulation and subcellular compartmentation. FEBS Letters 584: 2965–2973.

Nagegowda DA, Gupta P. 2020. Advances in biosynthesis, regulation, and metabolic engineering of plant specialized terpenoids. Plant Science 294: 110457.

Nagegowda DA, Gutensohn M, Wilkerson CG, Dudareva N. 2008. Two nearly identical terpene synthases catalyze the formation of nerolidol and linalool in snapdragon flowers. The Plant Journal 55: 224–239.

Ohara K. 2003. Limonene production in tobacco with Perilla limonene synthase cDNA. Journal of Experimental Botany 54: 2635–2642.

Parker GL, Smith LK, Baxendale IR. 2016. Development of the industrial synthesis of vitamin A. Tetrahedron 72: 1645–1652.

Philips M, Leon P, Boronat A, Rodriguezconcepcion M. 2008. The plastidial MEP pathway: unified nomenclature and resources. Trends in Plant Science 13: 619–623.

Purkayastha A, Mathur S, Verma V, Sharma S, Dasgupta I. 2010. Virus-induced gene silencing in rice using a vector derived from a DNA virus. Planta 232: 1531–1540.

Sato-Masumoto N, Ito M. 2014. Two types of alcohol dehydrogenase from Perilla can form citral and perillaldehyde. Phytochemistry 104: 12–20.

Sharmeen J, Mahomoodally F, Zengin G, Maggi F. 2021. Essential Oils as Natural Sources of Fragrance Compounds for Cosmetics and Cosmeceuticals. Molecules 26: 666.

Singh AK, Dwivedi V, Rai A, Pal S, Reddy SGE, Rao DKV, Shasany AK, Nagegowda DA. 2015. Virus-induced gene silencing of W ithania somnifera squalene synthase negatively regulates sterol and defence-related genes resulting in reduced withanolides and biotic stress tolerance. Plant Biotechnology Journal 13: 1287–1299.

Singh AK, Kumar SR, Dwivedi V, Rai A, Pal S, Shasany AK, Nagegowda DA. 2017. A WRKY transcription factor from Withania somnifera regulates triterpenoid withanolide accumulation and biotic stress tolerance through modulation of phytosterol and defense pathways. New Phytologist 215: 1115–1131.

Tamura K, Stecher G, Kumar S. 2021. MEGA11: Molecular Evolutionary Genetics Analysis Version 11 (FU Battistuzzi, Ed.). Molecular Biology and Evolution 38: 3022–3027.

Tan CS, Hassan M, Mohamed Hussein ZA, Ismail I, Ho KL, Ng CL, Zainal Z. 2018. Structural and kinetic studies of a novel nerol dehydrogenase from Persicaria minor, a nerol-specific enzyme for citral biosynthesis. Plant Physiology and Biochemistry 123: 359–368.

Thompson JD, Higgins DG, Gibson TJ. 1994. CLUSTAL W: improving the sensitivity of progressive multiple sequence alignment through sequence weighting, position-specific gap penalties and weight matrix choice. Nucleic Acids Research 22: 4673–4680.

Wu S, Schalk M, Clark A, Miles RB, Coates R, Chappell J. 2006. Redirection of cytosolic or plastidic isoprenoid precursors elevates terpene production in plants. Nature Biotechnology 24: 1441–1447.

Zhang X, Teixeira da Silva JA, Niu M, Zhang T, Liu H, Zheng F, Yuan Y, Li Y, Fang L, Zeng S, et al. 2021. Functional characterization of an Indian sandalwood (Santalum album L.) dual-localized bifunctional nerolidol/linalool synthase gene involved in stress response. Phytochemistry 183: 112610.

Zhang, Y., Zhang, Y., Tian, Q., Wei, L., Tang, W., Zhu, T., Zhou, Z., Wang, J., Xiao, H., Liu, Z. and Li, T., 2023. Dynamic floral scent profile of Epiphyllum oxypetalum and the cytosolic biosynthesis of trans-Geraniol through mevalonate pathway. bioRxiv, pp.2023–11.

Zhang J, Yu D, Zhang Y, Liu K, Xu K, Zhang F, Wang J, Tan G, Nie X, Ji Q, et al. 2017. Vacuum and Co-cultivation Agroinfiltration of (Germinated) Seeds Results in Tobacco Rattle Virus (TRV) Mediated Whole-Plant Virus-Induced Gene Silencing (VIGS) in Wheat and Maize. Frontiers in Plant Science 8.

Zhao Y, Chen Y, Gao M, Wang Y. 2024. Alcohol dehydrogenases regulated by a MYB44 transcription factor underlie Lauraceae citral biosynthesis. Plant Physiology 194: 1674–1691.

Zhou H, Wang S, Xie H-F, Liu G, Shamala LF, Pang J, Zhang Z, Ling T-J, Wei S. 2022. Cytosolic Nudix Hydrolase 1 Is Involved in Geranyl β-Primeveroside Production in Tea. Frontiers in Plant Science 13.

