## Supplemental Information for "Phylogenetically distant enzymes localized in cytosol and plastids drive citral biosynthesis in lemongrass"

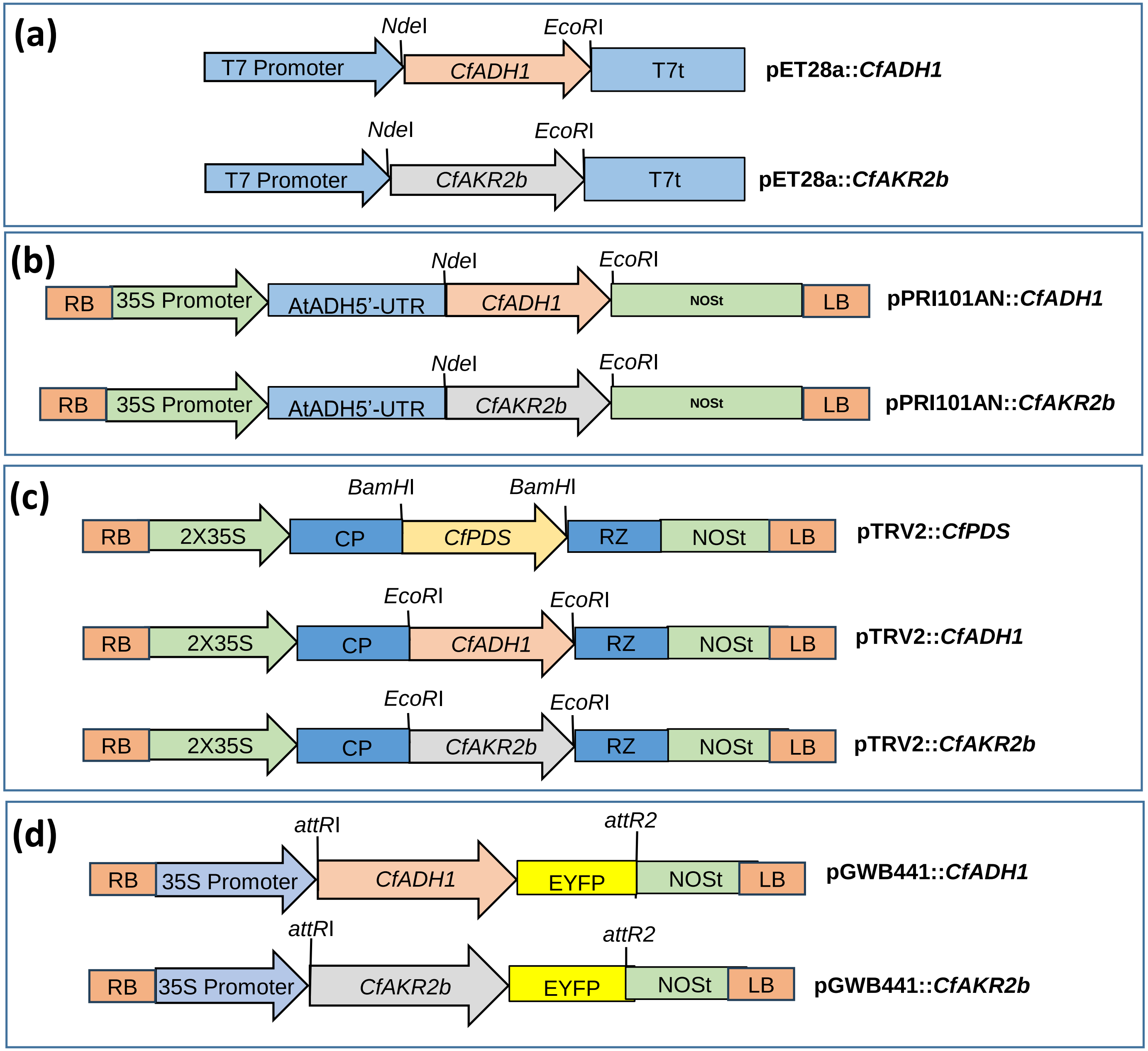

**Fig. S1** Schematic diagrams of constructs generated in this study. (a) pET28a-derived bacterial overexpression constructs. (b) pRI101AN-derived plant overexpression constructs. (c) pTRV2-derived VIGS constructs. (d) pGWB441-derived constructs used for subcellular localization. VIGS, virus induced gene silencing; AtADH, *Arabidopsis thaliana* alcohol dehydrogenase; NOST, Nopaline synthase terminator; UTR, untranslated region; CP, coat protein; RZ, self-cleaving ribozyme.

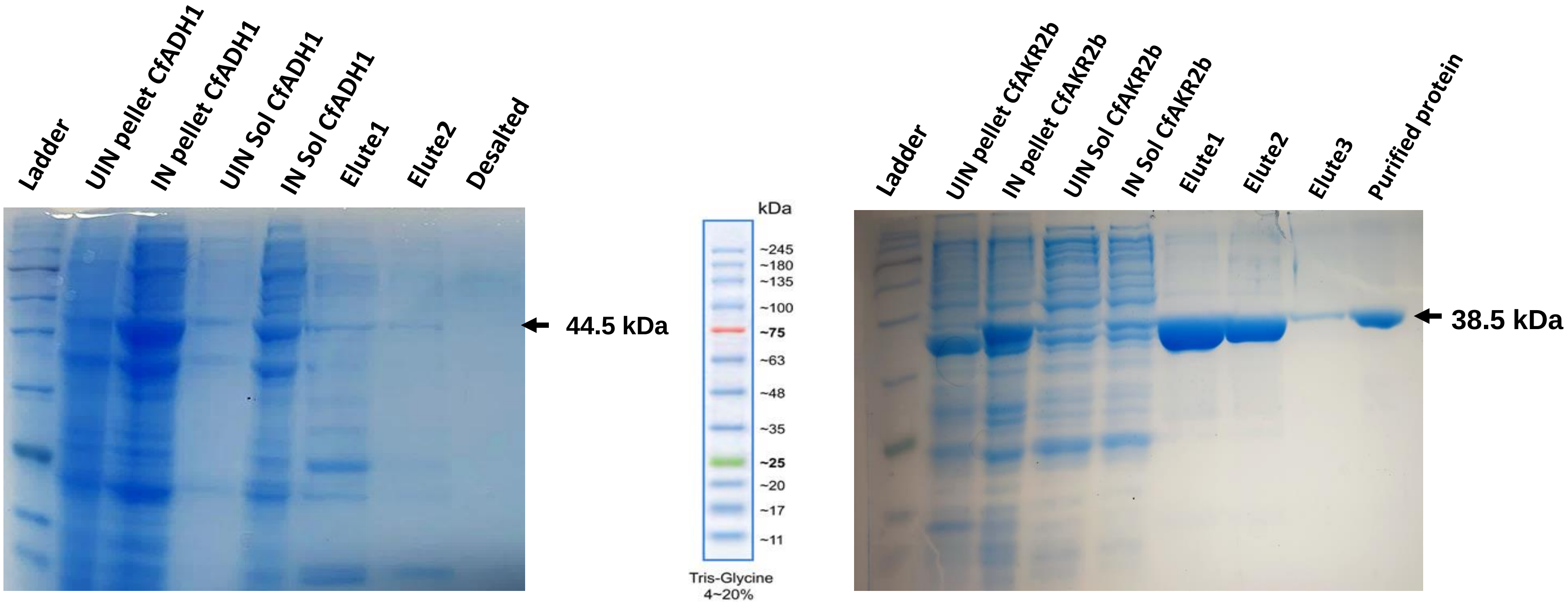

**Fig. S2** SDS-PAGE analysis of recombinant CfADH1 and CfAKR2b proteins expressed in *E. coli*.

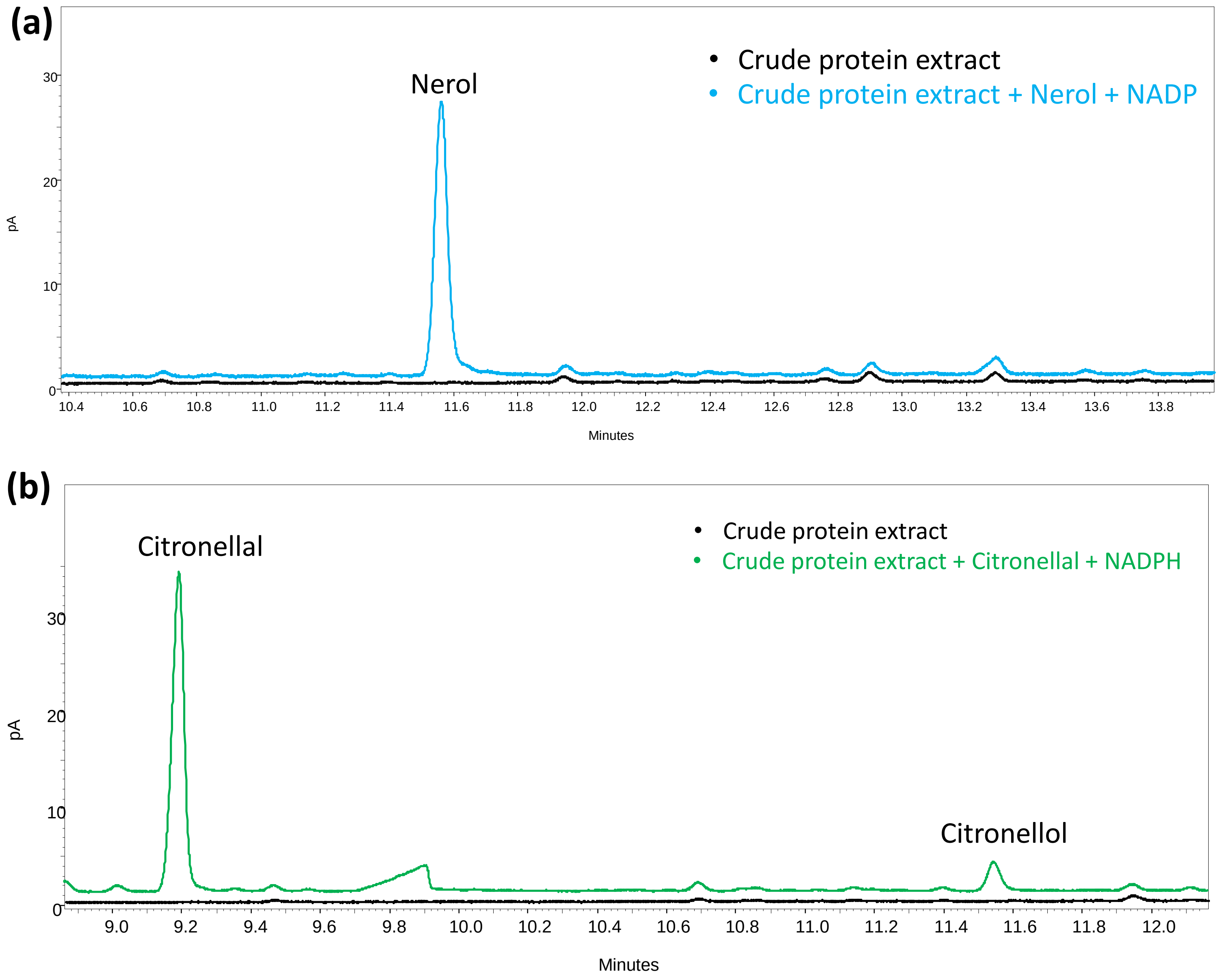

**Fig. S3** GC analysis of reaction products from assay consisting of lemongrass leaf crude protein with nerol and NADP cofactor (a), and with citronellal and NADPH cofactor (b). While there was no product formation in A, citronellol was formed from citronellal in (b).

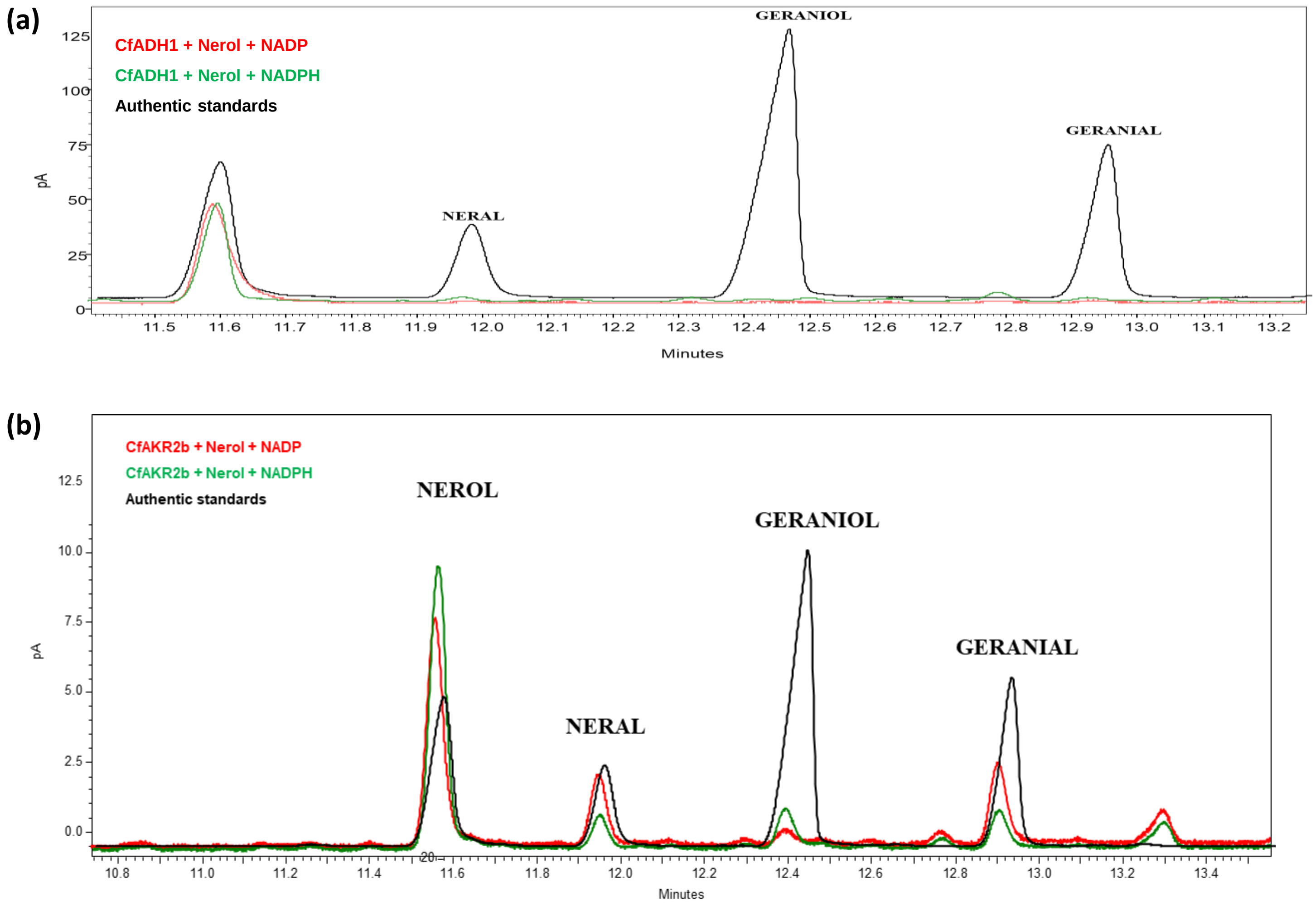

**Fig. S4.** GC analysis of reaction products from assay consisting of CfADH1 (a) and CfAKR2b (b) with nerol substrate and NADP or NADPH cofactor.

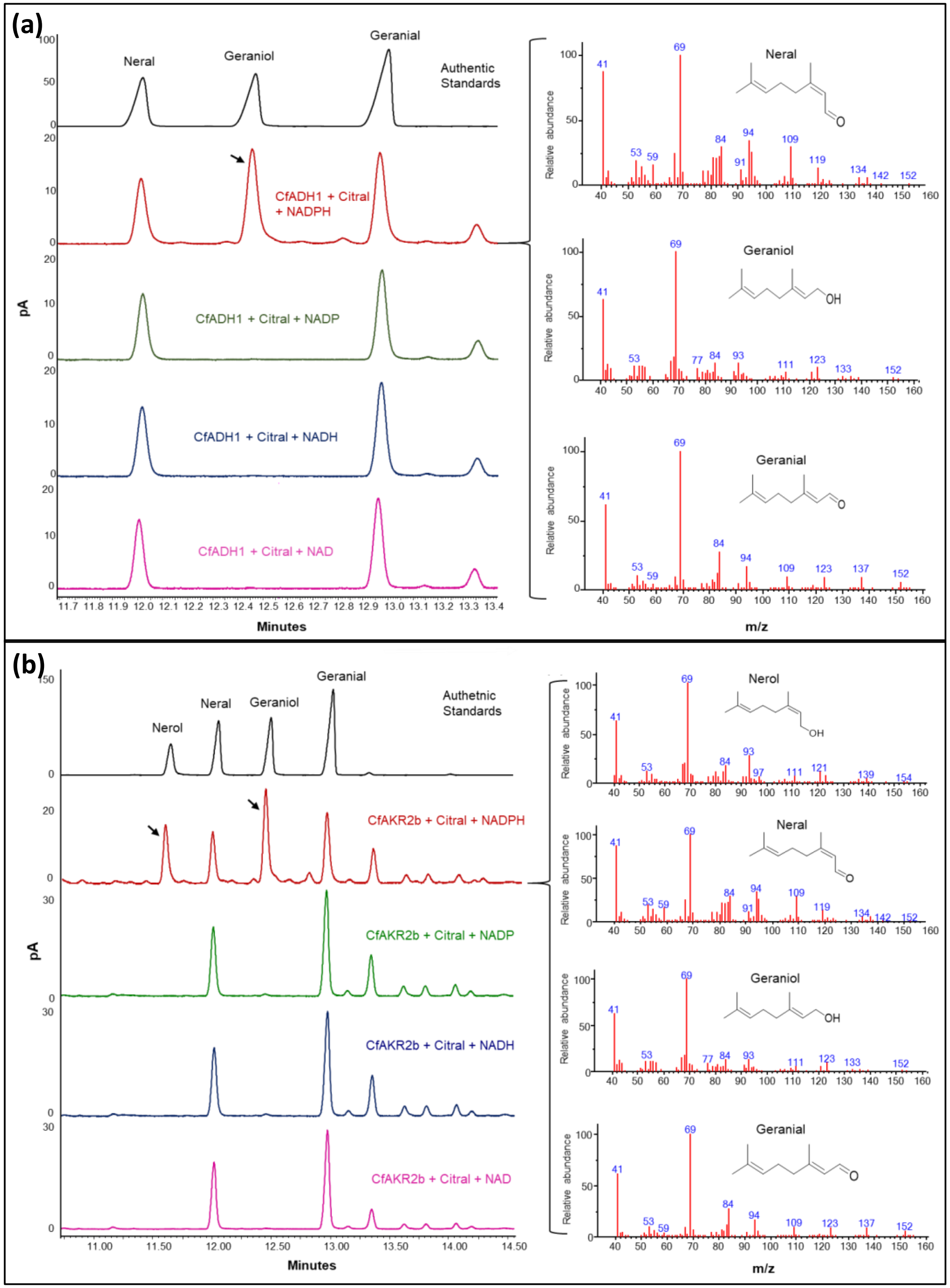

**Fig. S5** GC chromatograms of biochemical assay extracts of CfADH1 (a) and CfAKR2b (b) with citral and NAD or NADH or NADPH or NADP cofactors. The chromatograms are shown on the left and the GC-MS mass spectra of the products formed in CfADH1+citral+NADPH and CfAKRR2b+citral+NADPH assays are shown on the right.

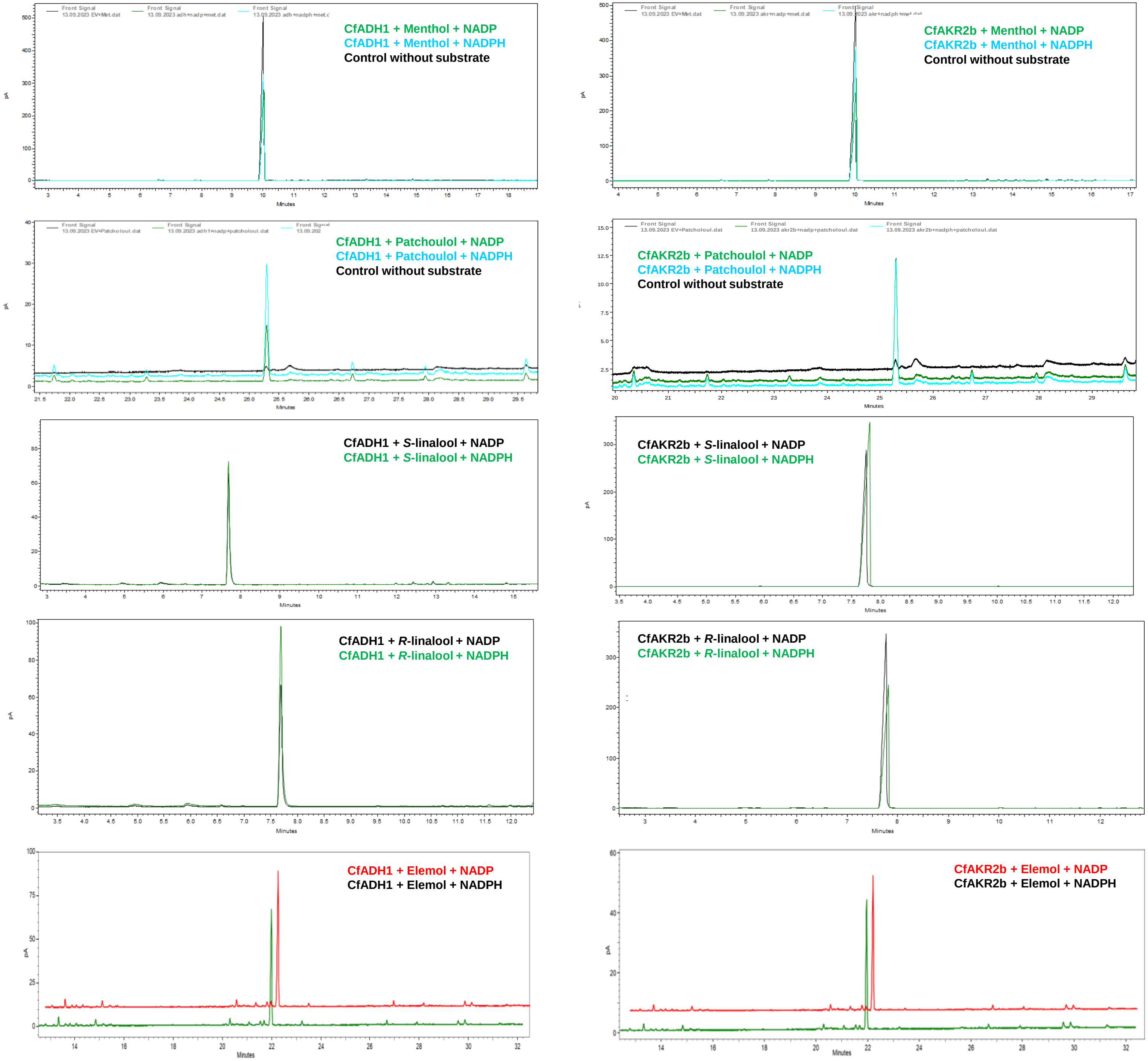

**Fig. S6.** GC analysis of reaction extracts from assay consisting of CfADH1 and CfAKR2b with different alcohol substrates using NADPH or NADP cofactors.

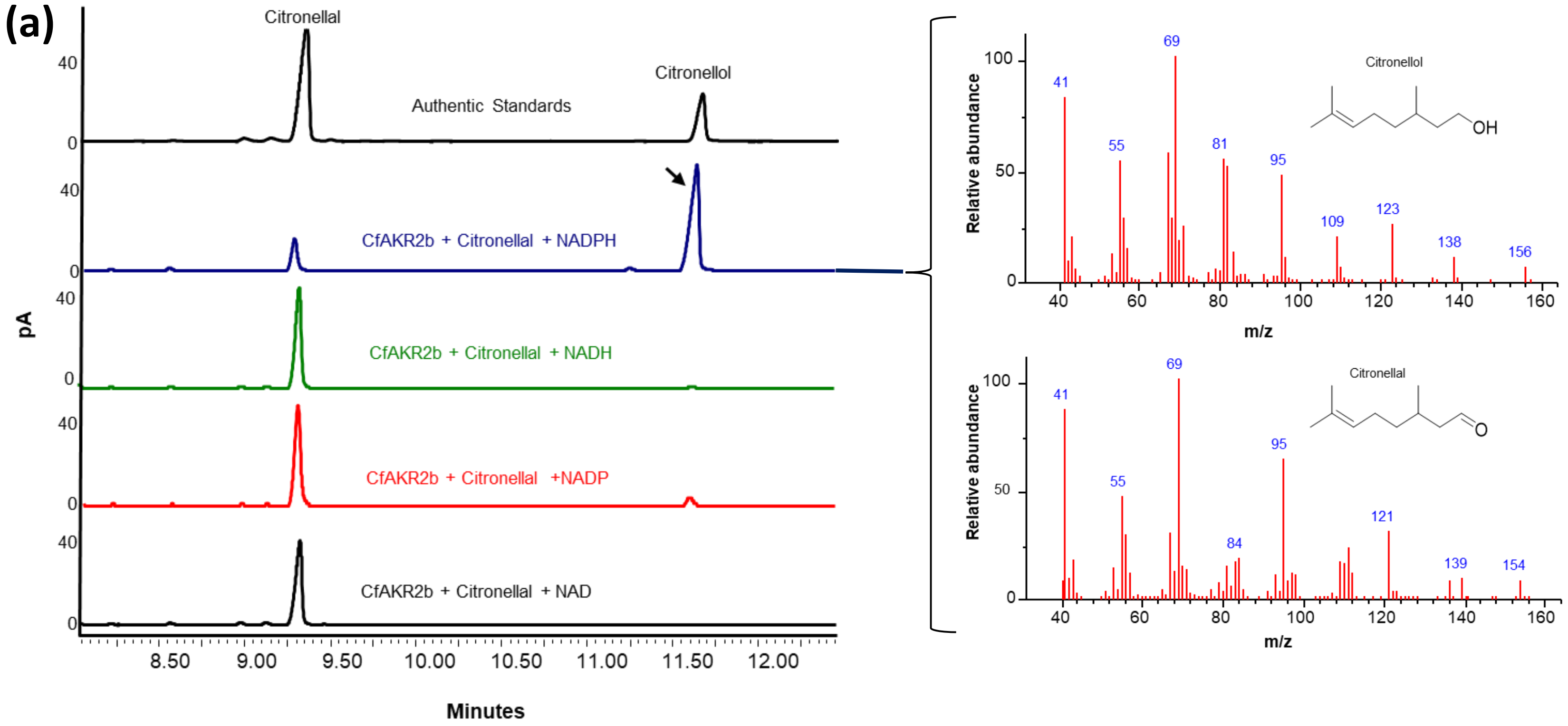

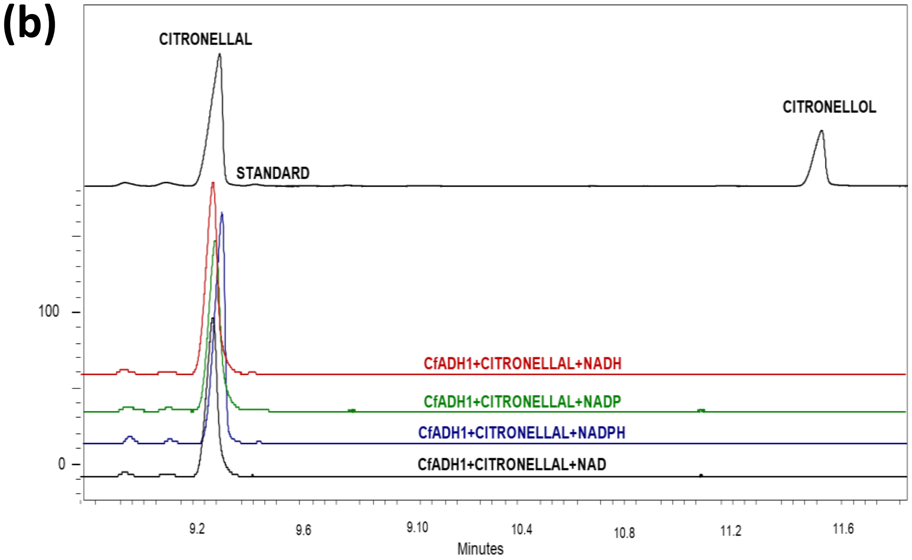

**Fig. S7** GC-MS analysis of reaction products from assay consisting of CfAKR2b (a) and CfADH1 (b) with citronellal substrate and different cofactors.

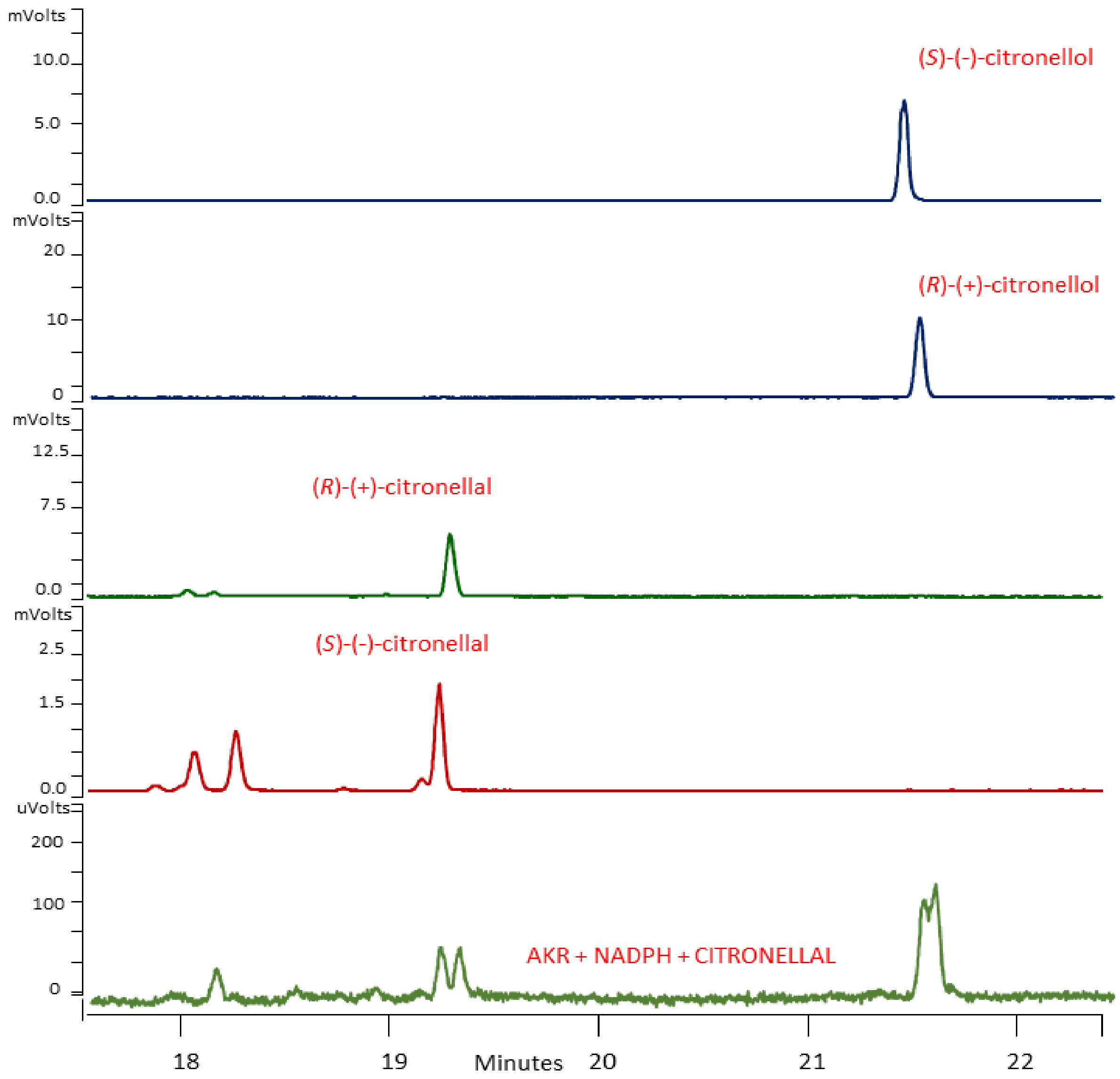

**Fig. S8.** Chiral GC analysis of biochemical assay products of CfAKR2b with *R* and *S* citronellal in presence of NADPH cofactor.

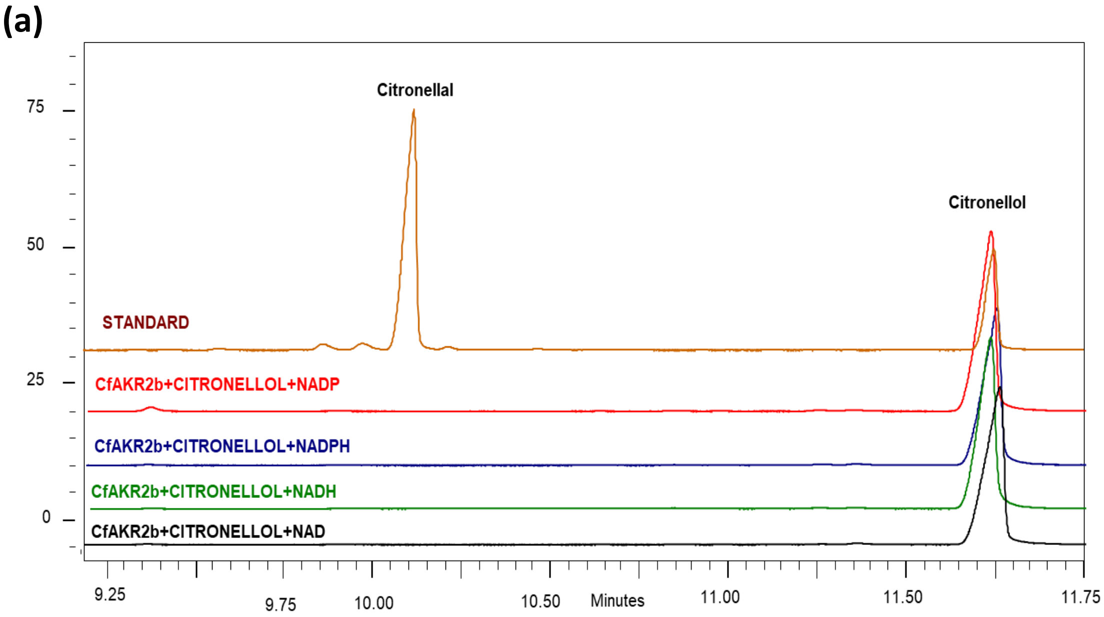

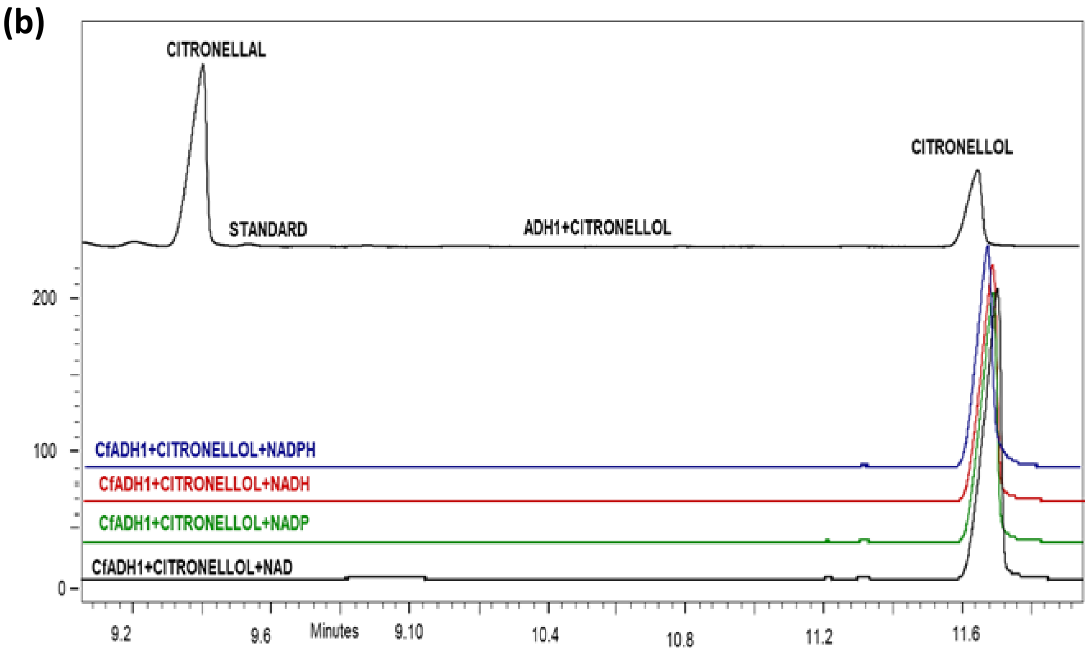

**Fig. S9** GC analysis of reaction extracts from assay consisting of CfADH1 (a) and CfAKR2b (b) with citronellol and NAD or NADP or NADH or NADP cofactors.

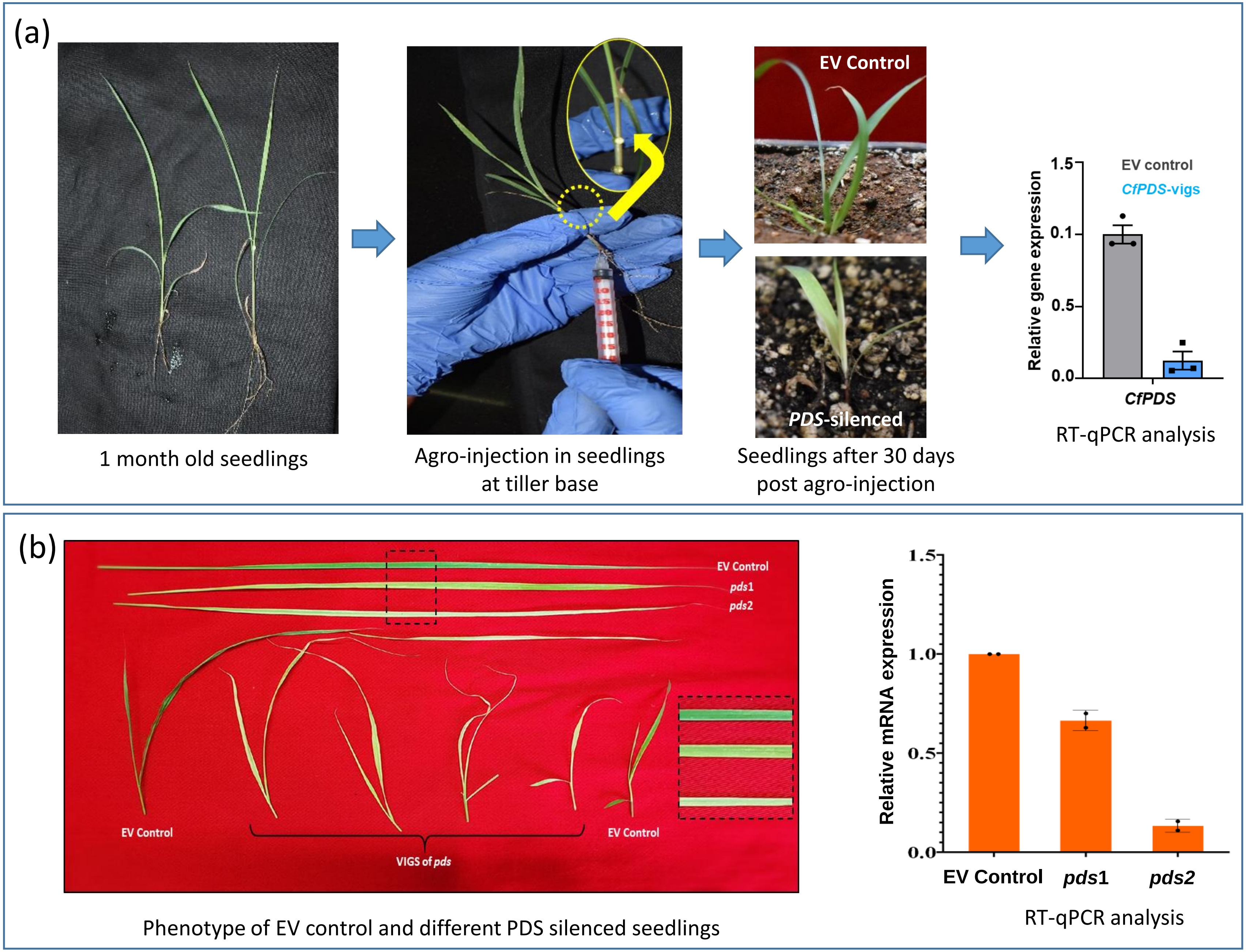

**Fig. S10** Schematics of virus-induced gene silencing (VIGS) methodology in lemongrass.

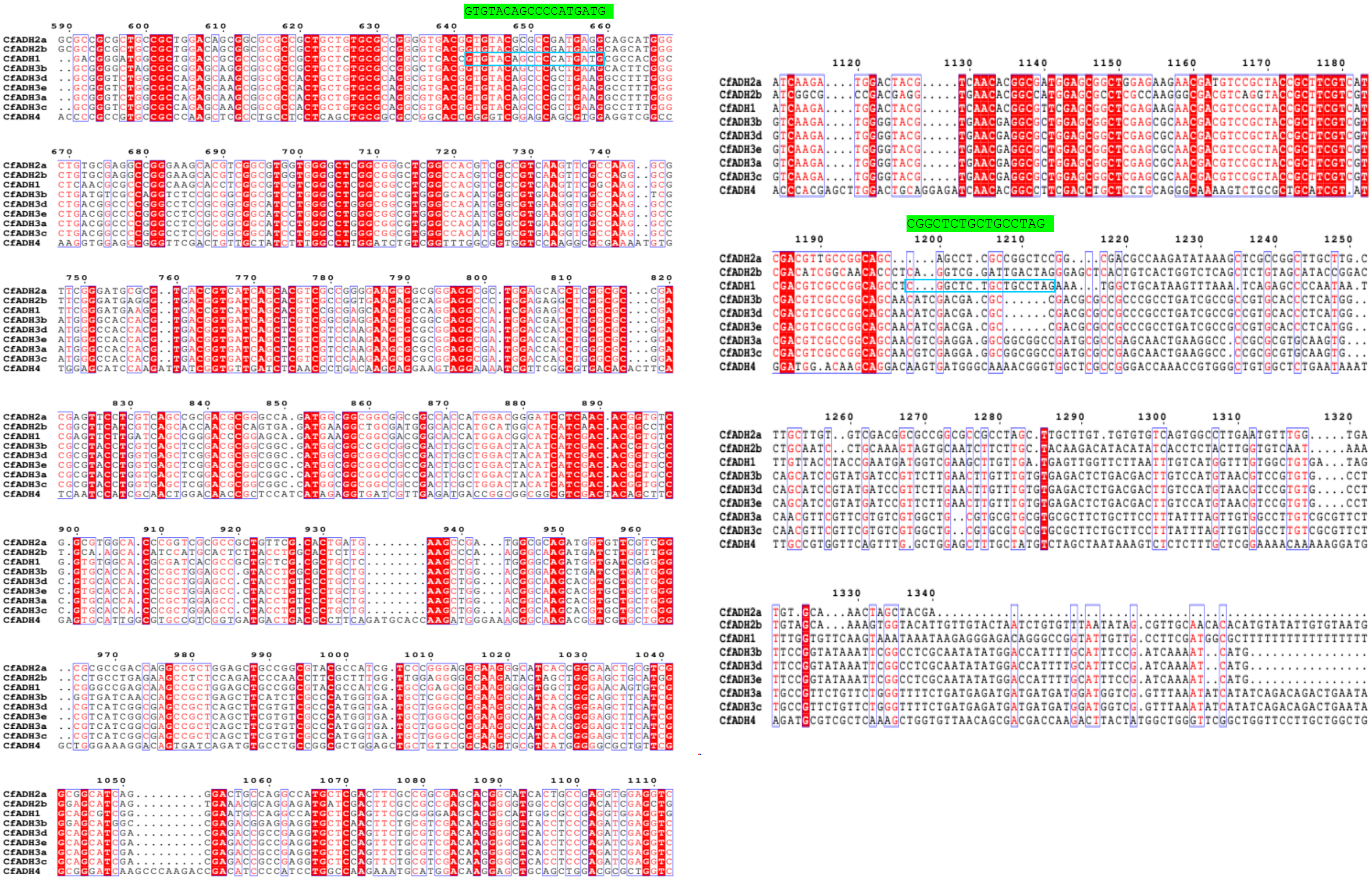

**Fig. S11** Multiple sequence alignment of DNA sequence of lemongrass CfADHs showing the location of CfADH1 VIGS primers. The CfADH1-specific primer region is boxed in blue and the corresponding nucleotide sequence is provided in green color.

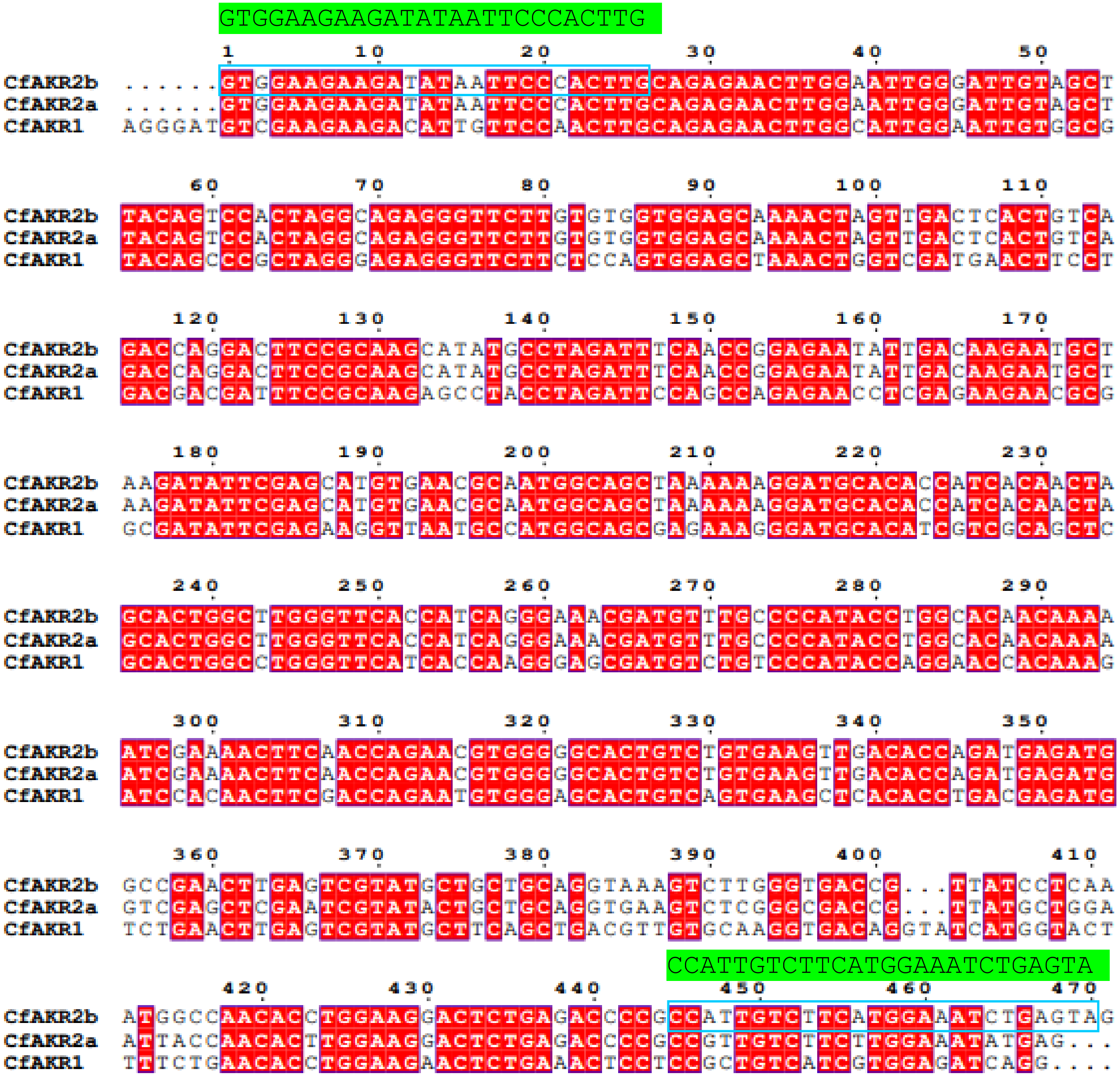

**Fig. S12** Multiple sequence alignment of DNA sequence of lemongrass CfAKRs showing the location of CfAKR2b VIGS primers. The CfAKR2b-specific primer region is boxed in blue and the corresponding nucleotide sequence is provided in green color.

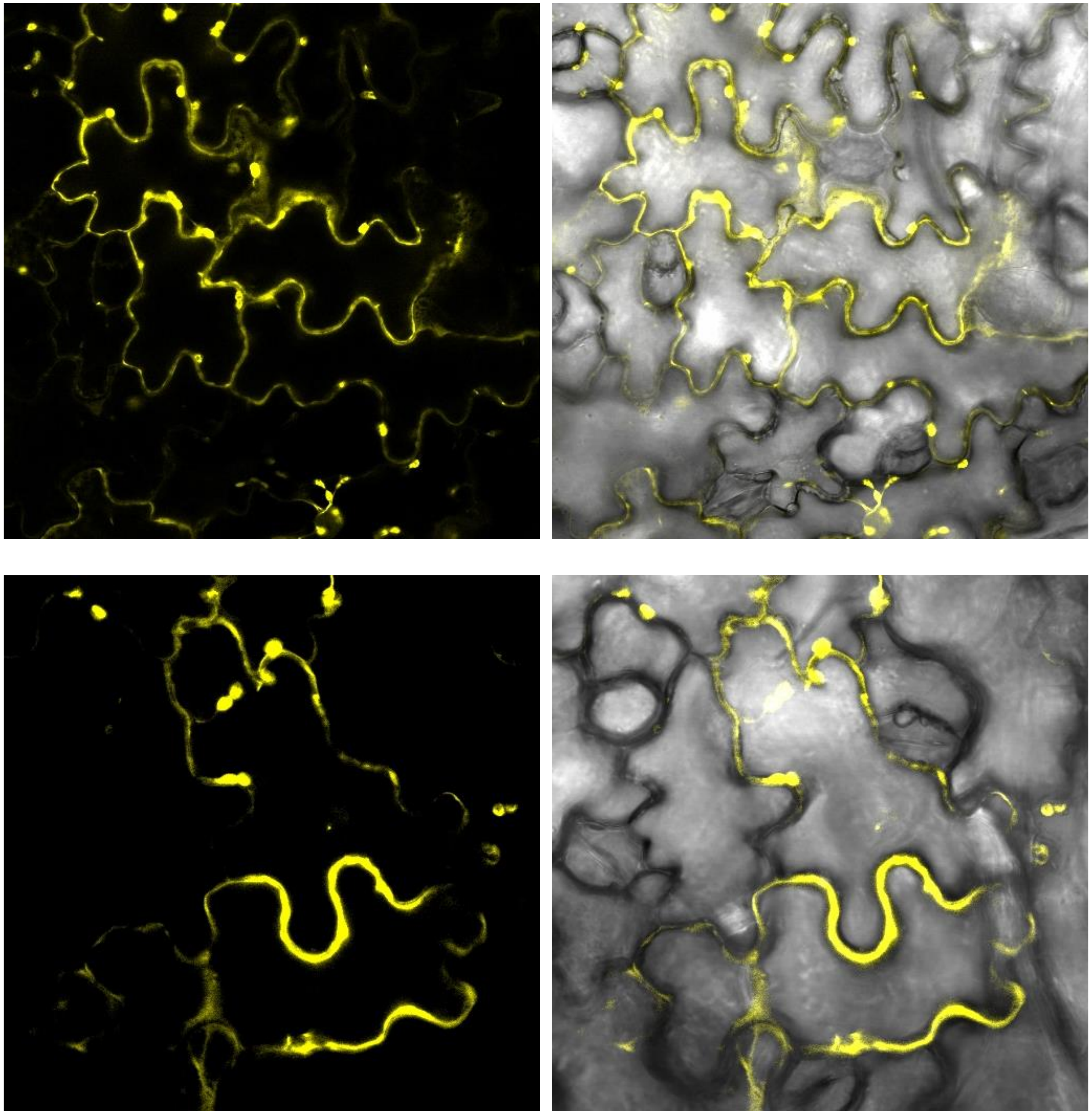

**Fig. S13** Additional confocal images showing the enlarged view of CfADH1-YFP expressing cells. Dual cytosolic and plastid localization is visible as diffused signal and as dots, respectively.

**(a)**

| | Lemon balm (Melissa officinalis) transcript ID | Protein length | Identity % | Positivity % | TPM |
|---|---|---|---|---|---|
| CfADH1 | TRINITY_DN4073_c0_g1_i19 | 360 | 60 | 75 | 51.91 |
| | TRINITY_DN22142_c0_g1_i1 | 360 | 57 | 73 | 85.17 |
| CfAKR2b | TRINITY_DN1370_c0_g1_i12 | 343 | 76 | 87 | 0.76 |
| | TRINITY_DN134_c0_g1_i2 | 342 | 73 | 85 | 0.31 |

| | Verbena officinalis transcript ID | Protein length (aa) | Identity % | Positivity % | TPM |
|---|---|---|---|---|---|
| CfADH1 | TRINITY_DN2684_c0_g1_i5.p1 | 360 | 51 | 71 | 32.87 |
| | TRINITY_DN2372_c0_g2_i3.p1 | 361 | 59 | 75 | 161.57 |
| | TRINITY_DN276_c2_g2_i1.p1 | 362 | 58 | 73 | 151.75 |
| CfAKR2b | TRINITY_DN569_c0_g1_i5.p1 | 343 | 75 | 86 | 157.13 |
| | TRINITY_DN4249_c1_g1_i1.p1 | 299 | 71 | 81 | 0.8 |

| | Litsea cubeba transcript ID | Protein length (aa) | Identity % | Positivity % | TPM |
|---|---|---|---|---|---|
| CfAKR2b | TRINITY_DN67_c0_g1_i15.p1 | 335 | 83 | 90 | 871.31 |
| | TRINITY_DN4926_c0_g1_i1 | 356 | 58 | 71 | 17.89 |

**(b)**

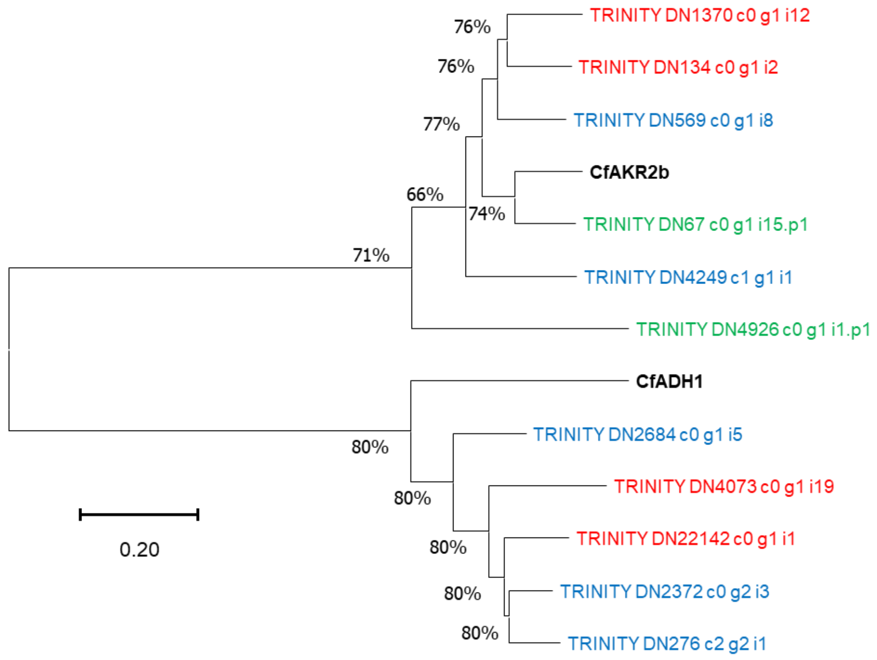

**Fig. S14** Identification of CfADH1 and CfAKR2b orthologs from lemon balm, *Verbena officinalis* and *Litsea cubeba*.

(a) Transcriptomes of *Melissa officinalis* (SRR5150719), *Verbena officinalis* (SRR19428485), and *Litsea cubeba* (SRR15882973) were retrieved from NCBI SRA database. They were assembled by Trinity and the assembled dataset was blasted with CfADH1 and CfAKR2b and the best hits were selected and represented in the table. Salman method of transcript abundance estimation was performed on trimmed raw data to get the TPM values of individual transcripts.

(b) Phylogenetic relationship of CfADH1 and CfAKR2b with homologs found in *Melissa officinalis* (red font), *Litsea cubeba* (green font) and *Verbena officinalis* (blue font). A phylogenetic tree was constructed by neighbour-joining method with bootstrap analysis from 1000 replicates using MEGA 11.0.11 (Tamura et al. 2021). The lengths of the branches show relative divergence among the reference CfADH1 and CfAKR2b amino acid sequences.

**Table S1.** List of oligonucleotide primers used in this study.

| List of Primers | | |
|---|---|---|
| Sl No | Gene | sequences |
| 1 | CfADH1_FL_F | CATATGATGTCATATCATTGTCGCGCG |
| 2 | CfADH1_FL_R | GAATTCCTAGGCAGCAGAGCCG |
| 3 | CfAKR2b_FL_F | GGATCCATGGCTGCCGCTTCC |
| 4 | CfAKR2b_FL_R | GAATTCCTACTCAGATTTCCATGAAGACAATGG |
| 5 | CfADH1_VIGS_F | GAATTCGTGTACAGCCCCATGATG |
| 6 | CfADH1_VIGS_R | GAATTCCTAGGCAGCAGAGCCG |
| 7 | CfAKR2b_VIGS_F | GAATTCGTGGAAGAAGATATAATTCCCACTTG |
| 8 | CfAKR2b_VIGS_R | GAATTCCTACTCAGATTTCCATGAAGACAATGG |
| 9 | CfPDS_F | GGATCCCACAATAAACTTTTTGGAAGCTGG |
| 10 | CfPDS_R | GGATCCCAGGAACACCCTGCTTTTTC |
| 11 | CfADH1_RT_F | CCTCTCCCCCTACAGCTTCTC |
| 12 | CfADH1_RT_R | GCCAGTCGTTCTTGATGATGTG |
| 13 | CfAKR2b_RT_F | GGGCATGTCCGCCTTCTAC |
| 14 | CfAKR2b_RT_R | CGTGGTGGATGAGCATGATC |
| 15 | CfPDS_RT_F | GCATTTTGATTGCTTTGAACAGA |
| 16 | CfPDS_RT_R | CCCTAGACCGAATGTGATCAACA |
| 17 | CfADH1_GW_F | GGGGACAAGTTTGTACAAAAAAGCAGGCTTAATGTCATATCATTGTCGCGCGG |
| 18 | CfADH1_GW_R | GGGGACCACTTTGTACAAGAAAGCTGGGTTGGCAGCAGAGCCGAGG |
| 19 | CfAKR2b_GE_F | GGGGACAAGTTTGTACAAAAAAGCAGGCTTAATGGCTGCCGCTTCCG |
| 20 | CfAKR2b_GW_R | GGGGACCACTTTGTACAAGAAAGCTGGGTTCTCAGATTTCCATGAAGACAATGGCG |

**Table S2** Accession numbers of characterized proteins used for phylogenetic analysis

| Protein name | Species | Accession number |
|---|---|---|
| LcADH28 | *Litsea cubeba* | **OR130230.1** |
| LcADH29 | *Litsea cubeba* | **OR130229.1** |
| CmADH | *Cinnamomum micranthum f. kanehirae* | **RWR86433.1** |
| Pm NeDH | *Persicaria minor* | **JX185716.1** |
| ZoGeDH | *Zingiber officinale* | **LC002206.1** |
| ObCAD | *Ocimum basilicum* | **AY879285.1** |
| ObGEDH | *Ocimum basilicum* | **AY879284.1** |
| PfAKR | *Perilla frutescens* | **JX629451** |
| PfGeDH | *Perilla frutescens* | **JX855836** |
| PcAKR | *Perilla citriodora* | **JX629452** |
| PcGeDH | *Perilla citriodora* | **JX855837** |
| PsAKR | *Perilla setoyensis* | **JX629453** |
| PsGeDH | *Perilla setoyensis* | **JX855838** |

**Table S3** *In silico* prediction of subcellular localization.

| Program | CfADH1 | CfAKR2b |
|---|---|---|
| iPSORT<br>https://ipsort.hgc.jp/ | Yes | Yes |
| WOLFPSORT<br>https://wolfpsort.hgc.jp/ | Chloroplast (14) | Cytosol (9) |
| Predotar<br>https://urgi.versailles.inra.fr/predotar/ | Possibly ER | None |
| TargetP 2.0<br>https://services.healthtech.dtu.dk/services/TargetP-2.0/ | Chloroplast (0.852) | Other (0.781) |
| Plant-mPLoc<br>http://www.csbio.sjtu.edu.cn/bioinf/plant-multi/ | Cytoplasm | Chloroplast |
| DeepLoc 2.0<br>https://services.healthtech.dtu.dk/services/DeepLoc-2.0/ | Chloroplast (0.5813)<br>Cytoplasm (0.4684 ) | Cytoplasm (0.8389) |
